# Estimating effective population size trajectories from time-series Identity-by-Descent (IBD) segments

**DOI:** 10.1101/2024.05.06.592728

**Authors:** Yilei Huang, Shai Carmi, Harald Ringbauer

**Affiliations:** Department of Archaeogenetics, Max Planck Institute for Evolutionary Anthropology, Leipzig, Germany; Bioinformatics Group, Institute of Computer Science, Universität Leipzig, Leipzig, Germany; Braun School of Public Health and Community Medicine, Hebrew University of Jerusalem, Jerusalem 9112102, Israel

## Abstract

Long, identical haplotypes shared between pairs of individuals, known as identity-by-descent (IBD) segments, result from recently shared co-ancestry. Various methods have been developed to utilize IBD sharing for demographic inference in contemporary DNA data. Recent methodological advances have enabled the screening for IBD in ancient DNA (aDNA) data, making demographic inference based on IBD also possible for aDNA. However, aDNA data typically have varying sampling times, but most demographic inference methods designed for modern data assume that sampling is contemporaneous. Here, we present TTNE (Time-Transect Ne), which models time-transect sampling to improve inference of recent effective population size trajectories. Using simulations, we show that utilizing IBD sharing in time series has increased resolution to infer recent fluctuations in effective population sizes compared to methods that only use contemporaneous samples. Finally, we developed an approach for estimating and modeling IBD detection errors in empirical IBD analysis. To showcase the practical utility of TTNE, we applied it to two time transects of ancient genomes, individuals associated with the Corded Ware Culture (CWC) and Medieval England. In both cases, we found evidence of a growing population, a signal consistent with archaeological records.

## Introduction

Published ancient DNA (aDNA) data has surged in recent years. In well-studied and well-preserved regions, this growing dataset now enables researchers to track demographic changes and evolution over time, which was previously only feasible for a few model organisms with short generation time [e.g. Burke et al., 2010, Bergland et al., 2014] or in rare cases, for well-documented wild populations in long-term field project [e.g. Chen et al., 2019, Lamichhaney et al., 2020]. Recent regional aDNA studies have generated time transect data, for example, in Cambridgeshire in the UK [Hui et al., 2024] and Denmark [Allentoft et al., 2024a].

Several computational methods were developed to infer the demographic history of a population, including its population size dynamics, based on genomic data from a sample of contemporary individuals [Browning and Browning, 2015, Fournier et al., 2023]. However, most of these methods cannot explicitly model time-series genomic data. Consequently, although many ancient DNA studies have time-series genomes from a single site or region, the analyses at the temporal dimension are often only descriptive, such as tracking ancestry compositions or allele frequency changes over time [e.g. Mathieson et al., 2015, Patterson et al., 2022, Gretzinger et al., 2022, Posth et al., 2023]. These approaches offer easily interpretable descriptions of the population and provide evidence of migration, admixture, or selection. However, these methods do not incorporate the time dimension as one of their model parameters and, therefore, do not fully utilize the temporal information to infer important population genetic parameters. Our method presented here is in line with several recent developments that directly model the time dimension to study important aspects of evolution [e.g. Joseph and Pe’er, 2019, Buffalo and Coop, 2020, Mathieson and Terhorst, 2022].

IBD segments are long, identical genomic regions shared by pairs of individuals, an unambiguous signal for recent genealogical connections as recombination events rapidly break them apart. Therefore, IBD sharing represents an ideal signal for investigating recent demography. Recent methodological advances have enabled robust IBD detection in large aDNA datasets (e.g., the tool ANCIBD [Ringbauer et al., 2023] or an alternative pipeline using IBDseq [Browning and Browning, 2013] as in Allentoft et al. [2024b]). Several methods have utilized IBD signal in modern populations to estimate various aspects of recent demography [e.g. Palamara et al., 2012, Palamara and Pe’er, 2013, Ralph and Coop, 2013, Browning and Browning, 2015, Al-Asadi et al., 2019, Nait Saada et al., 2020, Cai et al., 2023]. In addition, a recent method designed for aDNA, HapNe [Fournier et al., 2023], can estimate Ne from either IBD or linkage disequilibrium (LD). However, all these methods assume that all samples are contemporaneous or make approximations when samples are temporally stratified. As a result, if samples span a wide time interval, subtle bias may arise due to the simplifying assumption of time homogeneity.

Here, we present TTNE, a method to estimate recent effective population size (*N*_*e*_) trajectories from time-series aDNA data based on identity-by-descent (IBD) segments. Our method TTNE explicitly models IBD sharing in samples from different time points and directly utilizes this temporal dimension to improve the estimate of *N*_*e*_ trajectory. We evaluated our method using coalescent simulations, which show that it can accurately estimate recent *N*_*e*_ while avoiding overfitting using a regularization scheme. We compared the performance of IBD-based estimates of recent *N*_*e*_ under different sampling strategies, namely, all samples being contemporaneous vs. samples being temporally stratified. We demonstrate that the latter allows a substantially more accurate estimate of *N*_*e*_ when there is major demographic turnover over a relatively short period. Finally, we apply TTNE to individuals associated with Corded Ware (CW) culture and individuals in the British Isles from the Anglo-Saxon migration to the late Medieval period. We found evidence of population growth in both cases, consistent with archaeological and historical records.

## 1 Methods

Here, we describe our model for estimating population size trajectories using time-series IBD sharing. First, we will derive the theoretical IBD length distribution under a given demography. Our inference scheme then maximizes the composite likelihood of observed IBD length distributions with respect to demography.

Throughout, we assume a panmictic population with varying population sizes **N**_**e**_ over time. We use the bold font **N**_**e**_ to denote a vector of float-point values representing *N*_*e*_ at each generation backward in time. We assume that we have samples from *n* different time points and denote the set of samples in each time point *t*_*i*_ (measured by number of generations backward in time) by *S*_*i*_ with sample size *n*_*i*_. Moreover, we denote the set of sample sets by 𝒮={*s*_1_…*s*_n_}, and the set of sampling times by 𝒯 ={*t*_1_…*t*_n_} (Fig.1). Our model assumes that IBD segments longer than a given length cutoff *l* all have their most recent common ancestors younger than *G* generations before the oldest samples. This approximation has been used in several demographic inference schemes that utilize IBD sharing [e.g. Palamara et al., 2012, Browning and Browning, 2015]. Thus we aim to infer **N**_**e**_, a vector of length *T*_*max*_ = *G* + (max(𝒯) − min (𝒯)) .

**Figure 1:**
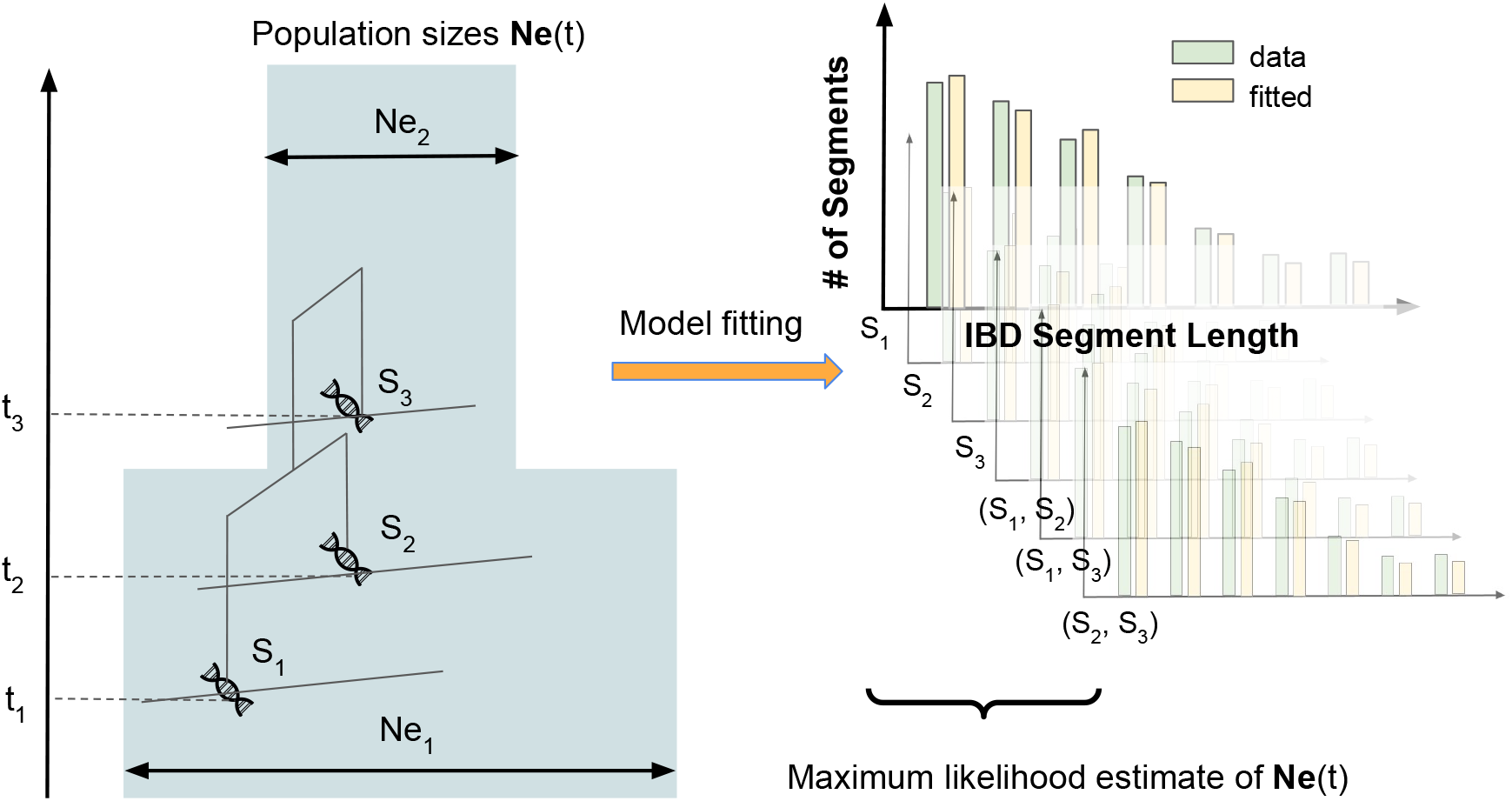
Schematics of using time-series IBD segments for demographic inference. A graphical representation of TTN_E_ model. The model assumes multiple sample sets *S*_*i*_ collected from different time points *t*_*i*_. TTN_E_ then fits a **N**_**_e_**_ vector to match the observed IBD sharing between samples, both within each sample set and across pairs of sample sets.

### 1.1 IBD distribution under a given demography

Throughout, we use the genetic map in Morgan to measure the lengths of IBD tracts, which, as a natural recombination distance measure, simplifies the calculations. We follow a long-standing analytical framework [Palamara et al., 2012, Palamara and Pe’er, 2013, Ralph and Coop, 2013, Browning and Browning, 2015], using the specific approach of [Ringbauer et al., 2017]. As explained there, the expected number of segments of length *l* originating from *t* generations backward in time, 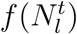, can be expressed as the product of two factors:

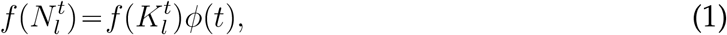

where the first factor 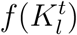 denotes the average number of uninterrupted ancestry blocks of length *l* at time *t* backward in time (which does not depend on the demography) and the second factor *ϕ*(*t*) the single-locus coalescent time density.

For any two haplotypes *h*_1_ ∈*S*_*i*_,*h*_2_∈*S*_*j*_, the time difference is Δ*t*=|*t*_*i −*_ *t*_*j*_| (Fig.2a), where *t*_*i*_,*t*_*j*_ are the times when the genomes *i,j* have lived. Assuming a Poisson model of recombination along a chromosome of length *L*, for *t >* max(*t*_*i*_,*t*_*j*_), one can modify Eq. 4 from Ringbauer et al. [2017], by updating that when the sampling times of two haplotypes are separated temporally by Δ*t*, the total branch length is 2*t* − Δ instead of 2*t*:

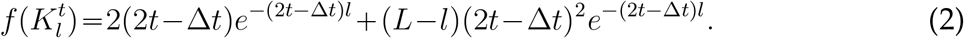

The first term describes blocks at one of the two ends of the chromosome, and the second describes blocks in the interior.

**Figure 2:**
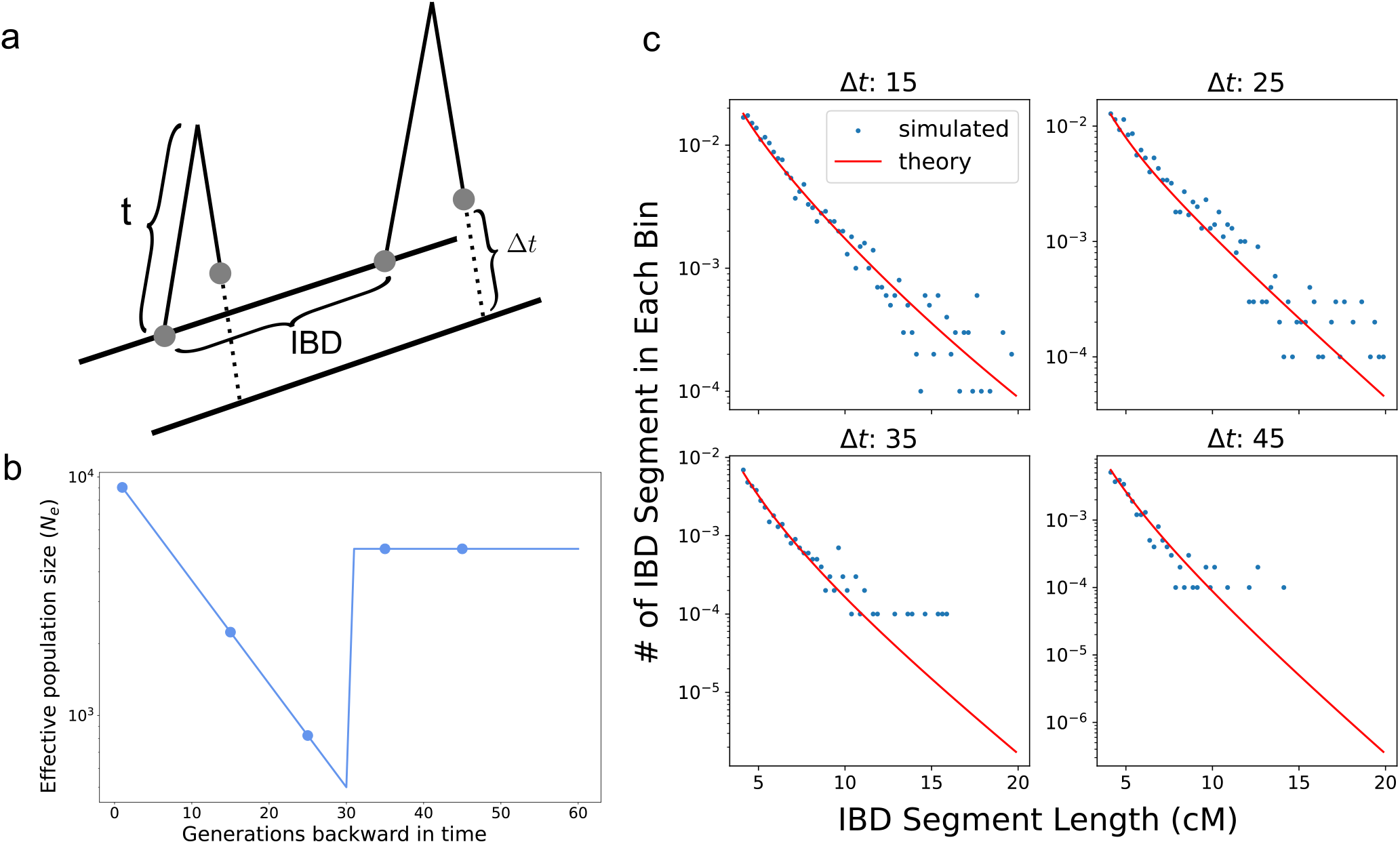
Haplotype sharing in temporally structured samples. **a** Schematic of sampling temporally structured haplotypes. The two sampled haplotypes are separated by Δ*t* generations. **b** We simulated a bottleneck demography to verify our model and sampled at *t* = 0,15,25,35,45. **c** For each pair of haplotypes, we took one at *t =* 0 and the other at various times in the past. We visualized the average number of IBD segments in each bin (bin size of 0.25cM, averaged over 10,000 replicates simulated with different random seeds) as blue dots and our model prediction as red lines.

For a single panmictic population with varying effective sizes **N**_**e**_, the single-locus coalescent rate *ϕ*(*t*) is

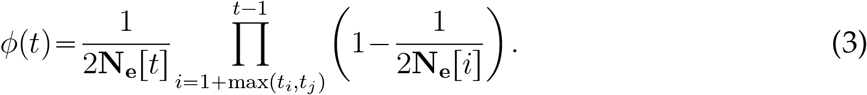

*l*

To obtain the number of segments of length *l* between the two haplotypes *h*_1_,*h*_2_, we sum up 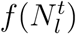 over *t* from max(*t*_*i*_,*t*_*j*_) to *G* generations into the past:

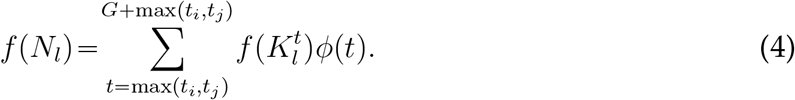

To verify this theoretical framework, we used MSPRIME [Baumdicker et al., 2022] to simulate chromosome 3 of a panmictic population with an instantaneous bottleneck at 30 generations backward in time, followed by exponential growth (illustrated in Fig.2b). We then obtained the empirical IBD sharing rate between a haplotype sampled at present (*t* = 0) and the others sampled at *t* = 15,25,35,45 using the ibd segment() function in tskit’s Python API. We compared this empirical rate with the theoretical prediction under the simulated demography, and we found that the empirical IBD rates closely match the theoretical predictions, indicating the accuracy of our framework (Fig.2c).

In the specific case of constant population size *N*, we can insert a continuous-time approximation into Eq. 3, in which *ϕ*(*t*) is an exponential random variable with rate 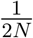. Integrating from Δ*t* to ∞ then yields a closed-form formula analogous to Eq. 4:

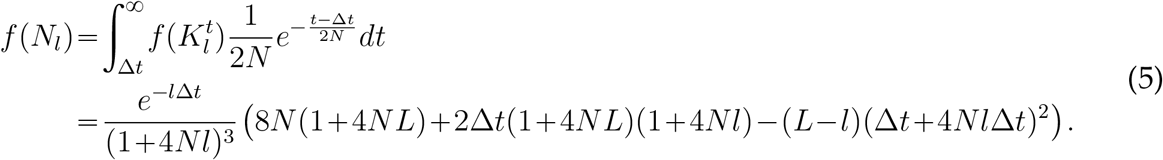

This equation is later used to estimate constant population size from the data to initialize our optimization subroutine.

### 1.2 Calculating the composite likelihood of observed IBD segments given a demography

As described in Ringbauer et al. [2017], the expected number of IBD segments whose length falls into a small length bin [*l,l* +Δ_*l*_] shared between a pair of haplotypes can be approximated by *f* (*N*_*l*_)Δ_*l*_. Within each length bin, we assume the number of segments follows a Poisson distribution with mean *n*_*hap*_*f* (*N*_*l*_)Δ_*l*_, where *n*_*hap*_ is the number of all possible haplotype pairs.

Assuming that each length bin is independent, the total likelihood is the product of Poisson likelihoods over all length bins. Denoting the log-likelihood of IBD segments shared between sample set *S*_*i*_ and *S*_*j*_ under a demography **N**_**e**_ by **ℒ**(*S*_*i*_,*S*_*j*_|**N**_**e**_), the composite likelihood of the observed IBD segments among all the sample sets *S* 𝒮 = {*S*_1_,*…*,*S*_*n*_} is:

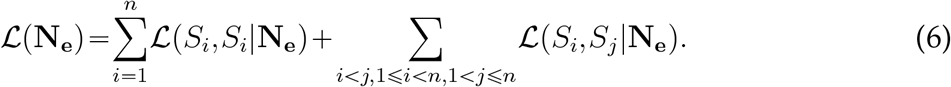

The first summation gives the likelihood of IBD sharing within each sampling cluster, while the second summation gives the likelihood of IBD sharing across all different sampling clusters. We note that the summands in Eq. 6 are not independent, as implicitly assumed when summing them to an overall likelihood. However, we show by simulations that this composite likelihood approximation works well in practice, and it has been used successfully in various other population genetic applications [e.g. Pritchard et al., 2000, Alexander et al., 2009, Ringbauer et al., 2017]

In practice, the inferred IBD segments are noisy estimates for two main reasons. First, IBD detection has imperfect power and precision, and the detected segments may have different lengths than the actual segments. Second, the set of observed IBD segments is only one random realization under a given demography. Therefore, we employed regularization to avoid over-fitting. Our regularization scheme is partly inspired by DeWitt et al. [2021], which uses ℓ_1_ norms of the first-order and third-order time derivatives of **N**_**e**_. Our early exploratory experiments showed that *ℓ*_1_ norm performed less well than *ℓ*_2_ norm for our optimization problems. Therefore, we use quadratic penalty functions.

Our penalty term consists of two parts. For the first term, we use the Hodrick–Prescott filter (H-P filter, [Hodrick and Prescott, 1997]), which is the sum of the square of the second difference,

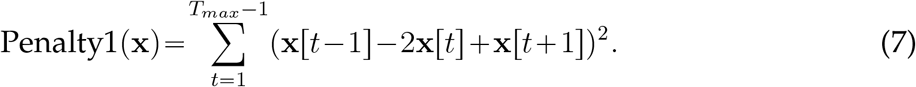

This term penalizes the curvature (the second derivative), effectively discouraging V or Λ-shaped solutions.

The second penalty term is the sum of the square of the first difference with Gaussian positional decay, which is related to the first derivative:

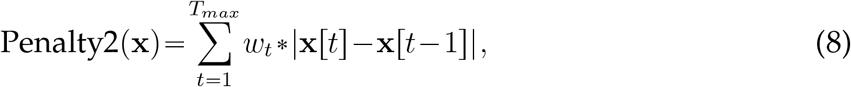

where

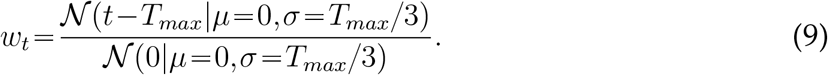

Qualitatively, this term penalizes changes in **N**_**e**_ deeper in time where coalescent signals from IBD diminish. We observed in our simulations that this term helps stabilize the *N*_*e*_ estimate at deeper time depths.

The full objective function of the optimization becomes:

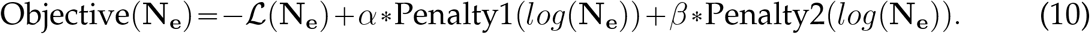

This regularization scheme is similar to several previous works [e.g. Ralph and Coop, 2013, DeWitt et al., 2021]. As in DeWitt et al. [2021], we apply a log transform before computing the penalty term because population size can vary over orders of magnitude during expansions and bottlenecks.

This regularization has two parameters, *α* and *β*, that determine the relative strength of the regularization terms. In practice, we found that different situations may entail different values of *α* to obtain a robust estimate of **N**_**e**_ (see Supp. Note S1 for how we adapt *α* automatically based on cross-validation). Unless otherwise stated, we fix *β* “250 as this term becomes negligible for the most recent part of inferred **N**_**e**_ and does not require fine-tuning for different scenarios.

### 1.3 Inference

We use the L-BFGS-B algorithm [Zhu et al., 1997] implemented in SciPy[Virtanen et al., 2020] to iteratively minimize the objective function Eq. 10 and to obtain the maximum likelihood estimator 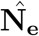 of the penalized composite likelihood,

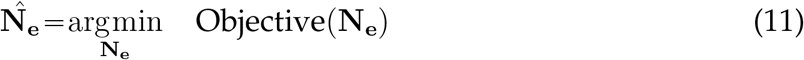

To initialize the optimization, we first obtain a maximum likelihood estimator 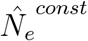 assuming a constant *N*_*e*_ using Eq.5. We then set the starting search value of **N**_**e**_ by 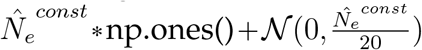. This small Gaussian noise helps the optimization algorithm explore a fuller search space.

To obtain confidence intervals, we bootstrap over chromosomes (sampling with replacement from the original 22 autosomes) 200 times and record the 2.5% and 97.5% percentile of each component of **N**_**e**_ as the confidence intervals.

### 1.4 Computing Posterior Distribution of TMRCA

After **N**_**e**_ is inferred, obtaining information about the time depth of which IBD segments are informative is helpful. Towards this end, we calculate the distribution of time to the most recent common ancestor (TMRCA) of an IBD segment. Given the **N**_**e**_, we can compute the posterior distribution of the TMRCA *T* of an observed IBD segment of length *l*. Applying Bayes’s theorem gives:

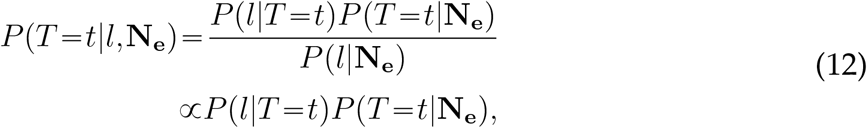

where *P* (*l*|*T* = *t*) denotes the probability density of having an IBD segment of length *l* spanning a chosen marker and originating from *t* generations ago, and it is given by Palamara et al. [2012], Baharian et al. [2016]:

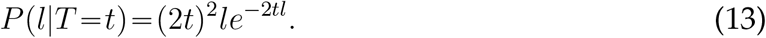

When Δ*t* generations separate the two samples, this term becomes

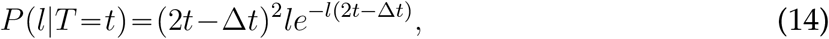

where *t* is measured from the younger sample. Finally, the second term *P* (*T* =*t*|**N**_**e**_) in Eg. 12 is given by Eq. 3. We verified this formula with simulations (Fig.S9). Our implementation of TTNE computes and visualizes this posterior distribution using the inferred 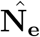 and its associated cumulative density function (see Fig.S7 for one example).

### 1.5 Modelling IBD Detection Error

Detecting IBD segments is a non-trivial task; thus, inferred IBD segments usually include errors such as false positives, imperfect recall, and length biases, which in turn affect the downstream demographic inference. To account for these errors, we use a model similar to Ralph and Coop [2013], Ringbauer et al. [2017] to incorporate three sources of errors. Denoting the theoretical rate of sharing (calculated from a given demographic history) an IBD block of length *y* by *λ*(*y*), then the observed rate of sharing 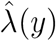 can be expressed as

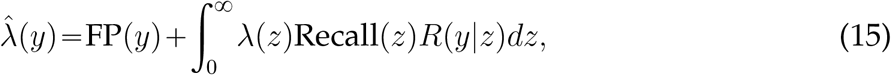

where FP(*y*) is the false positive rate of IBD block of length *y*, Recall(*z*) is the power to detect a segment of length *z* and *R*(*y*|*z*) is the probability that an IBD segment of true length *z* is detected being length *y*. See Supp. Note S2 for how we approximated these three error rates using estimates from simulated IBD segment data.

In practice, we limit the integral to an interval around *y*, as the inferred segment length is at most a few centimorgans away from the true length,

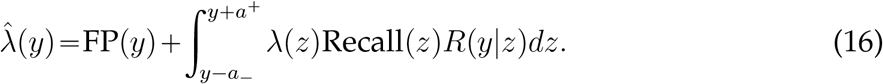

We chose *a*___ = *a*^+^ =5 in this study as we found in simulations that the inferred IBD length is typically within 5cM from its actual size (Fig.S3b,c). In our implementation, we evaluated this integral by summing over bins of width 0.25cM.

### 1.6 Simulating IBD segments

To generate simulated IBD segments, we used MSPRIME [Baumdicker et al., 2022] to simulate genome-wide genealogies and then extracted ground-truth IBD segments longer than 2cM using the ibd segments() function in the TreeSequence module provided in tskit’s Python API. Although we only use segments longer than 8cM for inference in our simulated data, we include shorter segments to simulate length bias errors (see 1.7). We simulated the 22 autosomes by stitching them together and used the HapMap [Gibbs et al., 2003] as the recombination map. The recombination rate between two consecutive chromosomes is set to log2 to simulate Mendelian inheritance logic, as described in https://tskit.dev/msprime/docs/stable/ancestry.html#multiple-chromosomes. We used the Wright-Fisher simulation engine of MSPRIME for the most recent 200 generations and then switched to the more efficient Hudson’s simulation engine. This hybrid simulation approach can more realistically capture IBD sharing among simulated samples [Nelson et al., 2020]. When simulating IBD segments, we terminated the ancestry simulation at 2500 generations backward in time, as deep coalescence effectively does not contribute to IBD sharing longer than a few centimorgans.

### 1.7 Simulating IBD Detection Errors

As explained in Section 1.5, we consider three types of IBD detection errors: false positives, recall, and length bias. To simulate false positives, we add segments to the ground-truth set. For example, if the false positive rate of IBD of length *l* is *f* (*l*), then we model the number of false positive segments whose length between *l* and *l* + Δ*l* as following a Poisson distribution with mean *f* (*l*)Δ*l*. We draw *n∼* Poisson(*f* (*l*)Δ*l*) segments of length 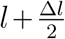 and add these false positives to the ground-truth set. In practice, we use Δ*l*=0.25*cM*, and we set the false positive rate to be equal to the expected sharing rate of a population with Ne=25,000 (Fig.4a). To simulate recall and length bias, for each segment in the ground-truth set, we keep it with the probability of recall(*l*). For simulation purposes, we use the functional form recall 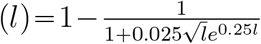, which is adapted from Ralph and Coop [2013] (Fig.4b). And if kept, the segment’s new length is its original length plus *ϵ*∼ 𝒩 (0,1.5). If a segment’s length drops below the length cutoff of 8cM (we use IBD segments longer than 8cM for inference in our simulation), this segment is discarded. Here, *ϵ* does not depend on *l* because we found a similar length bias across different segment lengths (Fig.S3b). An example of ground-truth IBD vs. ground-truth IBD with errors simulated as described above can be found in Fig.4c.

### 1.8 Empirical data analysis

We compiled published data from two periods and regions of interest to apply TTNE to empirical aDNA time-transect data. We selected a subset of unrelated individuals passing the coverage requirement for IBD calling (>1x for 1240k data and >0.25x for WGS data, [Ringbauer et al., 2023]). To filter ancestry outliers, we performed PCA using smartPCA. As widely done in ancient DNA analysis, we projected ancient genomes onto coordinates calculated from modern West-Eurasian Human Origins samples using shrinkage=YES. We estimated error parameters by simulating IBD under the average coverage of the selected samples (see Supp. Note S2). We used the median date of each individual’s radiocarbon or context date and grouped samples by five generations (Fig.S17), assuming a generation time of 29 years. Grouping samples by time is necessary because runtime scales quadratically with the number of distinct sampling points. We found that this level of grouping has negligible effects on the estimated **Ne** (Supp. Note S3).

## 2 Results

### 2.1 Performance on ground truth IBD

We simulated IBD under three different demographic scenarios: first, a constant population at *Ne*=25,000; second, a constant population undergoing an instantaneous 10-fold bottleneck (from 50,000 to 5000) 30 generations ago and then growing back exponentially to 100,000 at a steady rate; and, third, an exponentially increasing population at a constant rate starting from 50 generations ago (from 10,000 to 250,000). To mimic applications to aDNA data for which only long and medium-length IBD can be robustly detected, we used simulated IBD longer than 8cM.

As a baseline, we first only used contemporaneous samples (the most common setting where all samples are assumed to be taken at *t*=0) and compared TTNE with HapNe-IBD. We found that both methods perform similarly in this common setting (first row in Fig.3). At deeper time depth, HapNe-IBD estimated **N**_**e**_ tends to converge to the average **N**_**e**_ over time, while TTNE tends to converge to the estimated **N**_**e**_ value from the more recent time. These different behaviors mainly stem from different choices of regularization.

**Figure 3:**
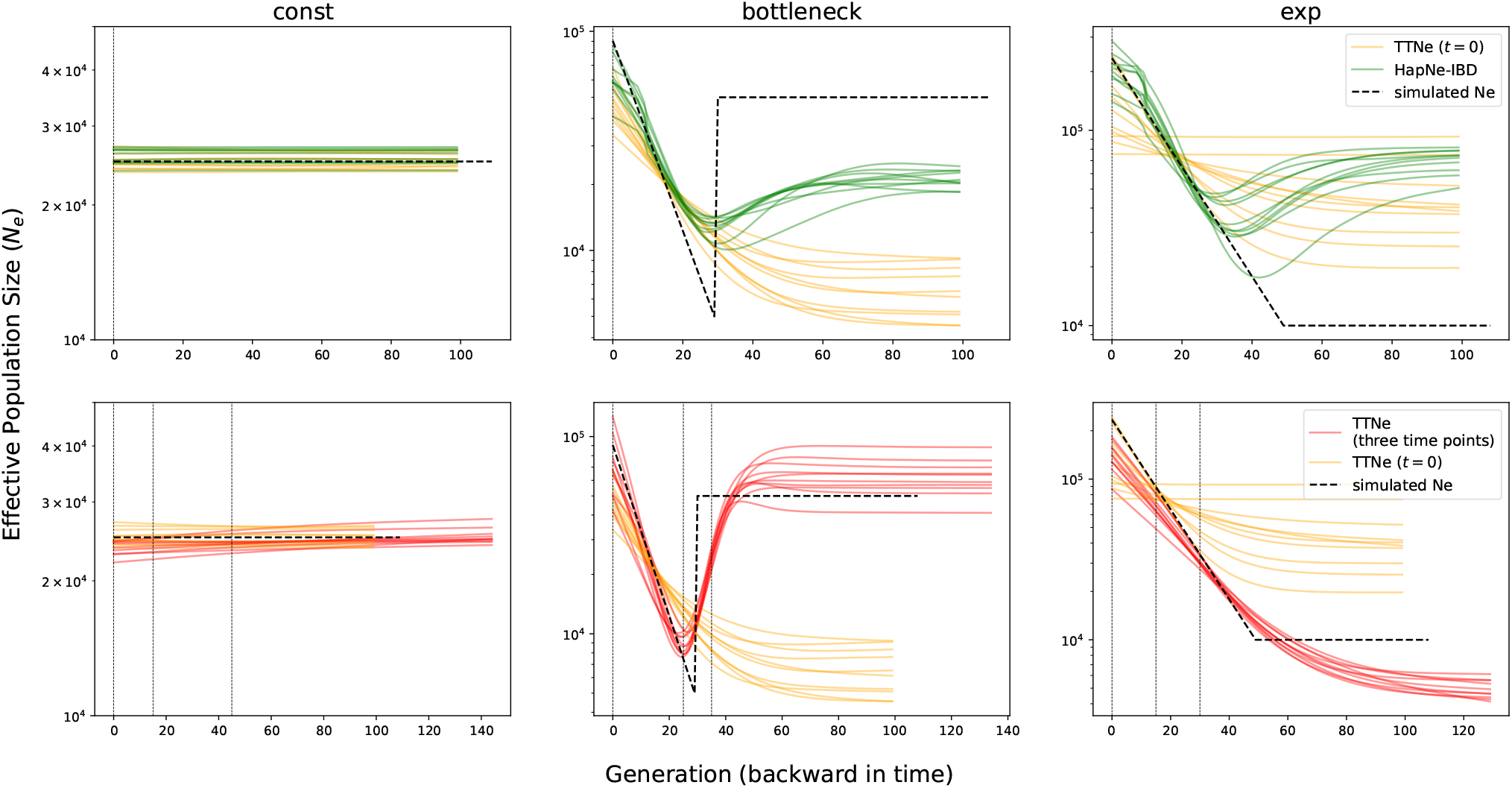
Performance of TTN_E_ in various simulated demographic scenarios. Each column visualizes one of the three simulated scenarios. The first row compares the results of TTN_E_ and HapNe-IBD using contemporaneous samples at *t =* 0. The second row visualizes the results of TTN_E_ using either contemporaneous samples at *t =* 0 (in orange) or samples from three different time points (in red, sampling times indicated by vertical lines). All results shown here use either *n=* 30 samples at each time point for simulations with three sampling times or *n =* 90 for simulations with only contemporaneous samples. We show ten independently simulated replicates for each scenario. The results of other sample sizes are shown in Fig.S11,S12,S13,S14,S15.

We simulated IBD sharing among three sampling points for each demographic scenario to evaluate the advantage of utilizing samples from multiple time points. For the constant demography, we sampled at *t*=0,15,45. For the bottleneck demography, we took samples at *t*=0,25,35; this sampling scheme has samples both before and after the bottleneck. For the exponential growth demography, we sampled at *t*=0,15,30. As no other method, to the best of our knowledge, can estimate **N**_**e**_ trajectories from time-series sampling, we could not directly compare TTNE to others in such sampling regimes. Instead, we compared the results of time-series samples against those of contemporaneous samples. To make the comparison fair, if *N* samples were taken at the three sampling time points, then 3*N* samples were taken at *t*=0 for the respective entirely contemporaneous sampling.

We found that time-series sampling recovers drastic changes in **N**_**e**_ more accurately than using only contemporaneous samples. For example, contemporary sampling cannot recover the large population size before the bottleneck in the bottleneck scenario. In contrast, the time-series sampling allows more accurate inference of the drastic changes of **Ne** around the bottleneck (second row in Fig.3). We also experimented with two sampling points where both sets of samples post-date the bottleneck. Even without samples predating the bottleneck, we can better recover both the magnitude of the bottleneck and the large population size before the bottleneck (Fig.S10), compared to the baseline with only contemporaneous samples. We also note that when the sample size becomes too small, both TTNE and HapNe-IBD lose the power to recover the true fluctuating **Ne** history and output **Ne** that is effectively constant over time (Fig.S11,S12). This flattening is expected in the low data regime when regularization dominates the signal.

#### 2.1.1 Performance on ground-truth IBD with simulated errors

We simulated IBD detection errors as described in 1.7 (also see Fig.4a,b,c). We then inferred **Ne** from the erroneous IBD call, applying the correction formula described in 1.5. We compared the inferred **Ne** with and without error correction (Fig.4d). We found that our correction formula effectively removes biases resulting from IBD detection errors. We note that in the simulation setting the error parameters (false positive rate, recall, and length bias) are correctly specified while, in practice, these parameters are unknown and can only be estimated; therefore, the estimated **Ne** may still suffer from some biases due to uncorrected misspecification.

**Figure 4:**
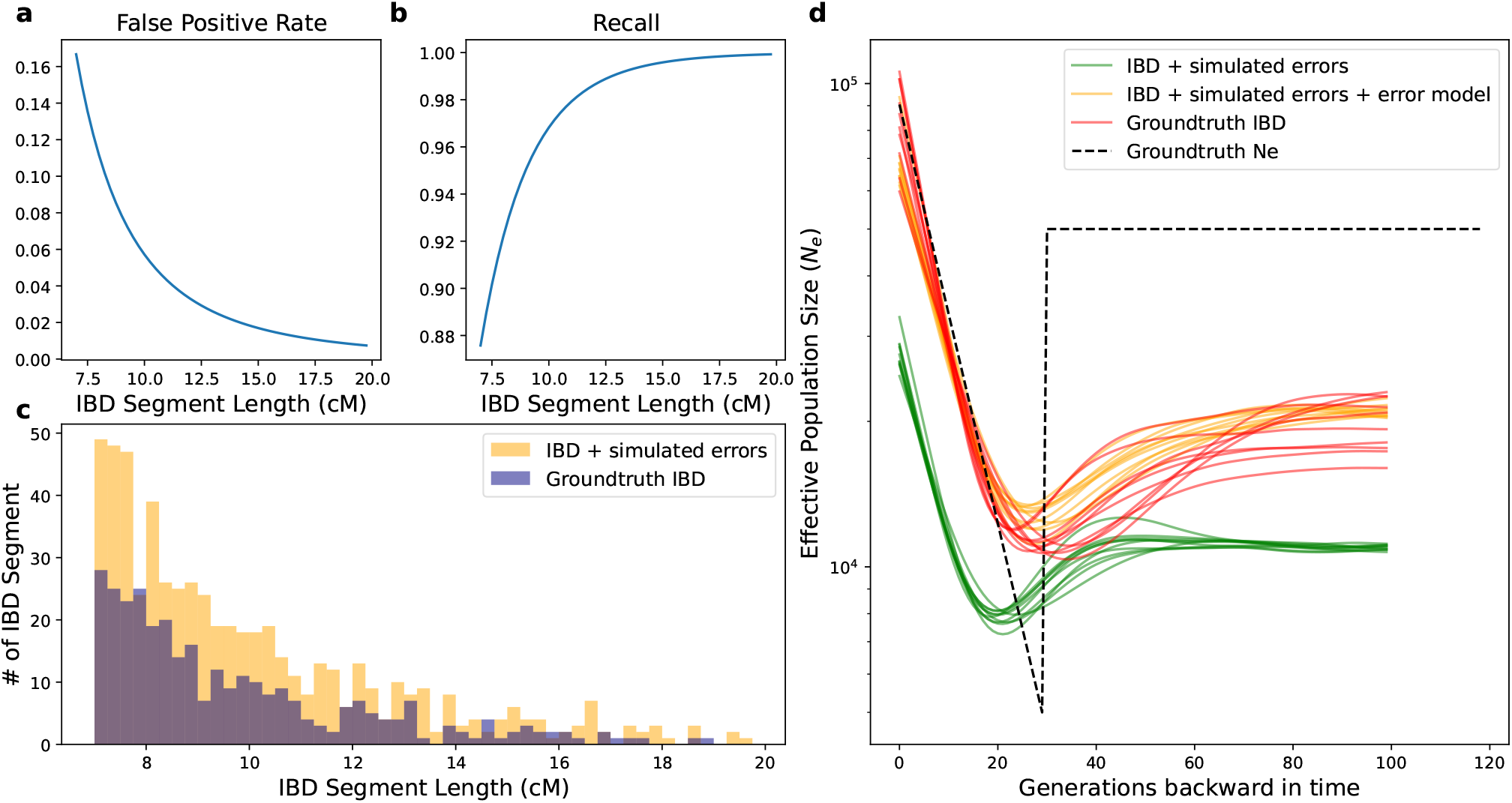
Explicit modeling alleviates biases from IBD detection errors. **a** Simulated false positive rate. We use the expected sharing rate of two random haplotypes drawn from a well-mixed population of effective size 25,000. In our simulated bottleneck scenario, the noise-to-signal ratio is then roughly 1:1. **b** Simulated recall. The recall function with respect to tract length *l* (in cM) is 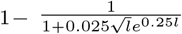. **c** Histogram of ground-truth IBD vs. IBD with simulated detection errors. The histogram visualizes IBD distribution for 180 diploid individuals sampled contemporaneously at *t =* 0 from the same bottleneck demography as in Fig.3. Simulated false positive and power are as in **a**,**b**, and the simulated length bias is drawn from a normal distribution with *μ =* 0,*σ =* 1.5 (so that 99.7% of the segments should be within 4.5cM from their original length). **d** Inferred *N*_*e*_ using ground-truth IBD (red), ground-truth IBD with simulated errors but without error correction (green), and with error correction (orange). 180 diploid individuals were sampled at *t =* 0. We show ten independent replicates for each scenario. In this simulation, the error model is specified as used in the simulation procedure.

### 2.2 Applications to individuals associated with the Corded Ware culture

Individuals associated with the Corded Ware culture, hereafter referred to as “CW,” were early carriers of steppe ancestry that arrived in a major genetic turnover across Europe in the third millennium BCE. CW individuals derive ca. ∼75% of their ancestry from a source very similar to the Pontic-Caspian steppe pastoralists[Haak et al., 2015, Allentoft et al., 2015, Papac et al., 2021], and the remaining ∼25% of their ancestry from a source similar to the earlier Chalcolithic farmers associated with Globular Amphora culture (GAC) [Ringbauer et al., 2023].

We imputed all the published Corded Ware genomes and ran ANCIBD using its default setting optimized for aDNA, as described in Ringbauer et al. [2023] (Supplementary Table 2). Because our demographic model assumes a single panmictic population, we used principle component analysis (PCA) to detect ancestry outliers (Fig.S6) and removed nine individuals as they fall outside the main cluster of other Corded Ware individuals (following a similar approach as in [Mathieson and Terhorst, 2022]). We only kept one individual with higher coverage from related pairs with over three IBD segments >12cM or the total sum of IBD segments > 12cM exceeding 100cM. Our filtering results in 42 samples suitable for IBD-based analysis. We grouped individuals of similar dates by five generations, resulting in four distinct groups with 12, 12, 8, and 8 individuals each (see Fig.S17 for sample date distribution. Each group’s median sampling time is depicted in Fig.5a) and excluded the remaining two individuals from *N*_*e*_ inference. We also explored different grouping (e.g., grouping by two or seven generations), which yielded qualitatively similar results (Fig S8).

**Figure 5:**
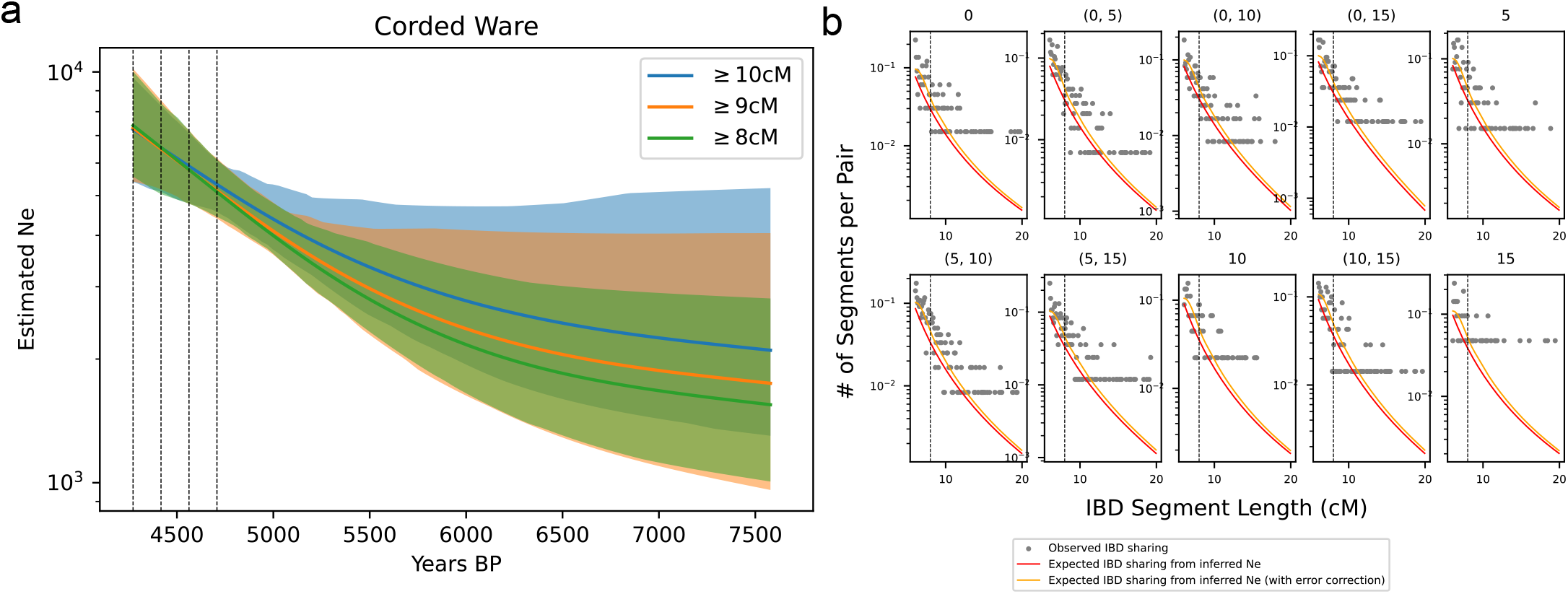
Inferred effective population size trajectory from Corded Ware genomes. **a** We visualize the estimated effective population size trajectories with different IBD length cutoffs over time. The shaded areas depict the 95% confidence intervals. The vertical dashed lines indicate the median age of samples in each of the four sampling clusters. The sample sizes at each sampling cluster are 12, 12, 8, and 8, respectively. **b** We visualize the fit between the empirically called IBD segments (gray dots) and the expected IBD sharing computed using the inferred effective population size trajectory (estimated with IBD length cutoff *≥*8cM, indicated by dashed vertical line). Detected IBD segments *≥*6cM are shown, although segments 6-8cM are not used for inference. The red and orange lines depict the expected IBD sharing assuming no detection errors and with detection errors, respectively. Four sampling clusters result in 4 + 4*3/2 = 10 IBD subplots, including both IBD sharing within one cluster and across two clusters (as described in Eq.6). The sub-figure title indicates the sampling time (measured in generations backward in time starting from the most recent sampling cluster) associated with the sample set. For example, 0 indicates that this subplot shows IBD within the sample set sampled at *t*=0. (0,5) indicates that this subplot depicts IBD between two sample sets, each dated to *t*= 0,*t*=5, respectively.

**Figure 6:**
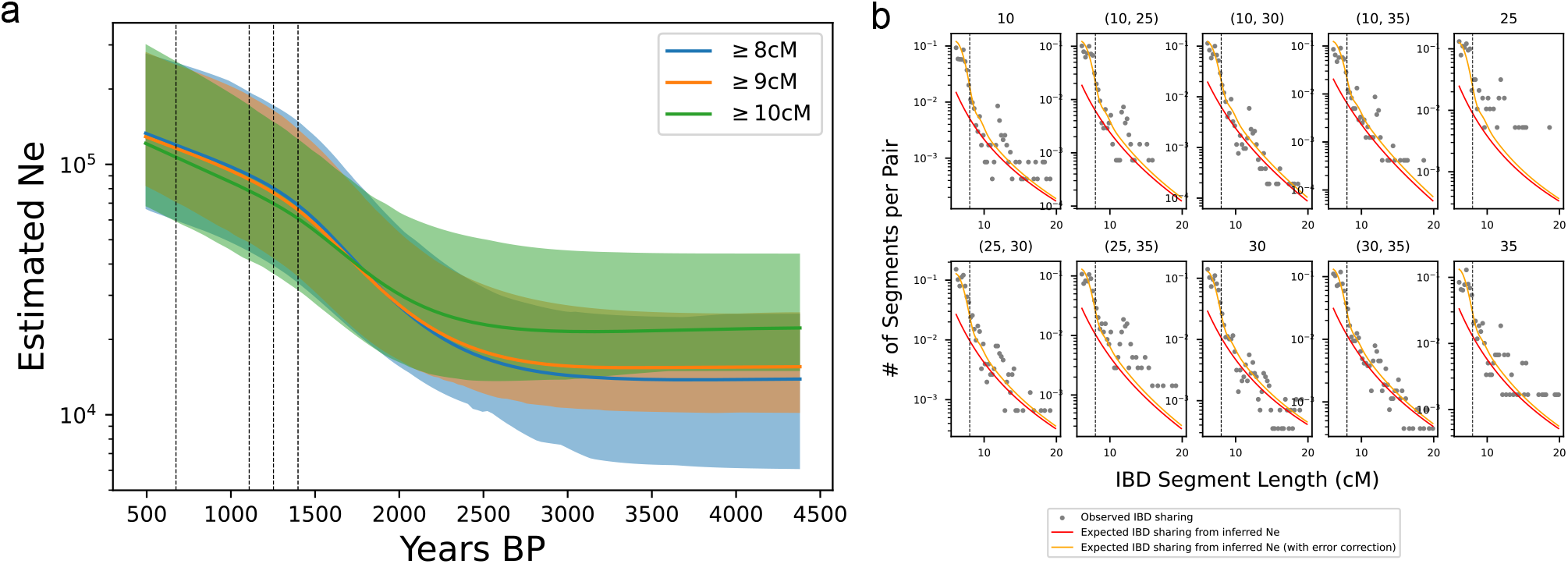
Inferred effective population size from British genomes. **a** Inferred **N**_**e**_ using different IBD length cutoffs. Shaded areas indicate 95% confidence intervals computed from bootstraps. We note that the inferred **N**_**e**_ starts 10 generations more recent than the median age of the youngest set of samples because we have used time-heterogeneity correction as described in the main text. **b** Similar to Fig.5, we visualize the fit between empirically called IBD segments *(≥*6cM) and the model predicted IBD using the **Ne** estimated using all segments *≥*8cM (indicated by vertical dashed black lines). The IBD segments between 6-8cM are not used during inference but are nonetheless visualized here for comparison.

We found a ∼3-fold population expansion in a ∼1000 year period starting from ∼5500BP (Fig.5). This signal is consistent with archaeological records that the CW individuals rapidly expanded into central, northern, and northeastern Europe from an unknown origin [Kristiansen et al., 2017]. Our inferred **Ne** curve does not provide a conclusive date on the onset of this population growth, possibly due to the relatively small sample size so that the regularization scheme pushes for a smoothed **Ne** solution. We also note that because the IBD signal diminishes exponentially with *t*, given that we can only infer comparably long IBD segments in imputed aDNA data, this fundamentally limits our ability to infer population sizes at deeper time depths. In Fig.S7, we visualized the cumulative density plot of the posterior distribution of TMRCA of IBD segments under this inferred CWC demography, showing that 97.5% IBD signals originate from coalescence within the past 50 generations. Therefore, the estimated **N**_**e**_ beyond 50 generations likely does not reflect the true effective population size, but rather is the result of regularization.

Finally, due to the admixture history of CW [**?**], LD-based methods to infer **Ne** may be biased by admixture LD. For instance, when applying hapNe-LD on published CW individuals, the software hapNe-LD suggests that there is substantial cross-chromosome LD and warns that the result may be substantially biased by recent admixture (Fig.S16). The inferred **Ne** either suggests a constant population when using early CWC individuals (dated to before 4460BP) or a recently declining population when using later CWC individuals (dated to after 4460BP). We note that admixture LD can lead to falsely inferred population contraction, as shown in Fournier et al. [2023].

### 2.3 Inferring population size trajectories in post-Anglo-Saxon Medieval Britain

The Anglo-Saxon migration in the 5-7th century CE constituted a significant gene flow into the British Isles ([Gretzinger et al., 2022]). We compiled ancient genomes from Gretzinger et al. [2022], which contains a large set of individuals collected from Southeastern Britain during the Anglo-Saxon migration period, and individuals from Hui et al. [2024], which includes samples collected from Cambridgeshire (also Southeastern Britain) during the Medieval age around the time of black death (1346-1353CE). The PCA (Fig.S18) suggests that genetic diversity within individuals from Hui et al. [2024] is a subset of that of individuals from Gretzinger et al. [2022], which is expected given that ancestry clines will stabilize and contract following a gene flow period. For data from Hui et al. [2024], we used individuals from sites labeled as “later medieval” and excluded sites labeled as “post-medieval” (1550-1855 CE) of which only seven samples remained after coverage filtering. The sites labeled as “later medieval” were dated from 940CE to 1561 CE, spanning approximately 20 generations. Because we found this level of sample time heterogeneity leads to a moderate upward bias in estimated **N**_**e**_ (Supp. Note S3) and because only a few samples are radiocarbon-dated, which prevents a more fine-grained time stratification, we group all samples from Hui et al. [2024] and apply the time heterogeneity correction formula described in Supp. Note S3. We use the median age of these samples (1239 CE) and set the time radius to be ten generations in both directions. For samples from Gretzinger et al. [2022], we used the median date of either radiocarbon or contextual dates for each sample and grouped samples of similar dates by five generations using a generation time of 29 years.

We inferred a stably growing effective population size from the early Middle Ages to the onset of the black death in England in the 14th Century. The inferred Medieval population growth from 500 to 1300 is consistent with historical estimates [Cipolla, 1972]. Agriculture expanded, and advancements in medieval technology allowed more land to be farmed. Population growth only stopped with the 14th-century Crisis of the Late Middle Ages (including famines and the Black Death in 1348) when population counts fell abruptly [Cipolla, 1972]. However, given that our latest samples are from the 1500s, the population decline is not yet observed in our data.

Notably, the inferred effective population size in Medieval Britain is an order of magnitude larger than the one in the Early Bronze Age Corded Ware (∼10^5^ compared to ∼ 10^4^), indicating much higher population densities in the Middle Ages. Therefore, our application to this period serves as a proof-of-concept use case showing that our tool can also infer effective population sizes in large Medieval populations with comparably low rates of IBD sharing - given that aDNA sample sizes are sufficiently large.

## 3 Discussion

We have introduced a tool for estimating population size trajectory from IBD segments in time-series ancient DNA data. We show using simulations that our method’s ability to utilize IBD sharing signals among multiple time points recovers population size changes more accurately than previous methods that assume that all samples are contemporaneous or fit an average **N**_**e**_ curve for time heterogeneous samples (e.g., HapNe-IBD). Unlike allele-frequency-based methods, IBD-based methods have a unique advantage because IBD-sharing signals originate exclusively in the most recent past (usually within 50-100 generations, or equivalently, ∼3000 years). The shallow time depth of IBD sharing shields this signal from confounding by deep population history and admixtures common in the human past that would bias allele-frequency-based methods. Moreover, using IBD-based methods is much more robust to ascertainment biases, as calling IBD works for various sets of ascertained variants and can even tolerate substantial genotyping errors as is typical in ancient DNA data [Ringbauer et al., 2023].

A limitation of our model is its assumption of a single, well-mixed population. Violations of this assumption complicate interpretations of the inferred changes in **N**_**e**_. As effective population size as estimated by our method is effectively the inverse of the coalescent rate, any factor that affects the coalescent rate could lead to changes in estimated **N**_**e**_ trajectory, including effects from migration, selection [Schrider et al., 2016, Cousins et al., 2024] and population structure [Mazet et al., 2015]). This limitation is not unique to TTNE, and any method that relies on the assumption of a panmictic population is similarly prone to misinterpretation [e.g. Browning and Browning, 2015, Mathieson and Terhorst, 2022, Fournier et al., 2023]. Therefore, one should always consider other factors that lead to changes in coalescent rate when interpreting the estimated **N**_**e**_ in

To showcase TTNE’s utility to infer effective population size trajectories in actual aDNA data, we applied it to two empirical aDNA datasets: individuals associated with the Corded Ware culture ca. 5000 years ago and individuals from post Anglo-Saxon Medieval Britain. In both cases, we found evidence of recent population growth. One caveat of our analysis of the British dataset is that only a relatively small fraction of individuals’ birth dates in Hui et al. [2024] can be unambiguously dated to pre- or post-black death (ca. 1350 CE). Therefore, we have grouped all later medieval samples together. Grouping individuals born before and after the black death together may lead to an underestimation of **N**_**e**_ in the most recent generations. To accurately estimate the regional demographic impact of black death, further studies could aim to generate data from dated individuals to construct a time-series dataset spanning the entire black death period.

As in-solution capture kits similar to the 1240k capture widely used in aDNA studies become widely available [Rohland et al., 2022] and whole genome sequencing is becoming more cost-effective, we anticipate that more studies will be able to generate genome-wide ancient DNA data at a large scale. As the published aDNA records keep growing rapidly, we anticipate that more studies will be able to utilize time-transect to study fine-scaled demographic changes in specific regions over time. Therefore, TTNE will also become relevant for currently less well-studied regions, helping to reveal past population size trajectories around the globe, particularly for periods where little direct historical evidence on past population sizes is available.

## 4 Data Availability

No new DNA data was generated for this study. The BAM files of individuals associated with the Corded Ware culture used in this study were downloaded from ENA repositories associated with their respective publications (Supplementary Table 4). The BAM files of individuals from Britain are either from Gretzinger et al. [2022] (ENA accession number PRJEB54899) or Hui et al. [2024] (ENA accession number PRJEB59976).

## 5 Code Availability

The development code of TTNE is deposited at https://github.com/hyl317/IBDTimeSeries. TTNE is available as a python package (https://pypi.org/project/TTNe/0.0.1a0/) and can be installed via pip.

## Author Contributions

We annotate author contributions using the CRediT Taxonomy labels (https://casrai.org/credit/). Where both authors serve in the same role, the degree of contribution is specified as ‘lead’, ‘equal’, or ‘support’.

- Conceptualization (Design of study) – lead: YH, HR; support: SC
- Software – YH; support: HR
- Formal Analysis – YH
- Data Curation – YH
- Visualization – YH
- Writing (original draft preparation) – lead: YH; support: HR
- Writing (review and editing) – YH, HR, SC
- Supervision – HR, SC

