## Supplementary Materials for "Estimating effective population size trajectories from time-series Identity-by-Descent (IBD) segments"

May 6, 2024

#### Supp. Note S1 Choosing regularization parameters

Our regularization has two components: the first is related to the second derivative of $\mathbf{N_e}$, and the second to the first derivative (Eq.7 and Eq.9 in main). The two components have weight $\alpha,\beta$ respectively. As stated in the main text, we chose to fix $\beta = 250$ throughout, as the second term is intended to stabilize the estimated Ne at deeper time ranges where there is little coalescence from long IBD segments. It only has very small effects on estimated recent $\mathbf{N_e}$ due to the positional decay we use.

However, we observed that choosing a single $\alpha$ that works across a wide range of demographics and sampling strategies was challenging. Therefore, we used 5-fold cross-validation to select the appropriate value for $\alpha$. We equally divide all pairs of individuals into five parts. We hold out one part for validation and use the remaining four parts of pairs to estimate $\mathbf{N_e}$ at a given $\alpha$. We note that pairs of individuals are not independent (as individuals are shared among pairs), violating the assumption for cross-validation. Therefore, we experimented with partitioning by chromosomes or individuals. However, we found that in both cases, the small sample sizes lead to noisy estimates.

We use the inferred $\mathbf{N_e}$ to calculate the so-called deviance statistics on the held-out validation set. The deviance statistics is a commonly used goodness-of-fit measure in Poisson regression. It is defined as follows,

$$D = 2\sum_{i=1}^{n}\left(y_i \log\left(\frac{y_i}{\lambda_i}\right) - (y_i - \lambda_i)\right),$$

where the summation is over small length bins, $\lambda_i$ denotes the number of IBD within that length bin predicted from the inferred $\mathbf{N_e}$, and $y_i$ the number of observed IBD segments falling within that length bin. The deviance statistic measures how the observed counts deviate from those predicted by the fitted model. It equals zero exactly when $y_i = \lambda_i \forall i$. When $y_i = 0$, we set the individual summand, otherwise undefined due to the log, to 0 by convention because $\lim_{x\to 0^+} x \log x = 0$.

We then use the average value of deviance statistics over the five validation sets, denoted by $D(\alpha)$, as the metric to select $\alpha$. We perform a grid search for $\alpha$ from 50 to 1e7, using 30 values evenly spaced on a log scale. Usually, one would choose the $\alpha$ that yields the smallest $D(\alpha)$. However, rapid oscillations of $\mathbf{N_e}$ will not substantially worsen the fit, even though that makes the $\mathbf{N_e}$ highly irregular. Based on this heuristic rationale, we aim to choose the strongest regularization (i.e., the smoothest predicted $\mathbf{N_e}$) that does not yield a substantially worse fit: We chose the biggest $\alpha$ so that $D(\alpha)$ remains within one plus the smallest $D(\alpha)$. Formally, the regularization $\alpha^*$ is then determined by

$$\alpha^* = \max\{\alpha : D(\alpha) \leqslant 1 + \min_{\alpha} D(\alpha)\}$$

We empirically observed that all other things being equal, a dataset with a bigger sample size tends to have a smaller $\alpha^*$, hence weaker regularization (Fig.S1). Heuristically, this is a desirable feature as more data contains more signal to infer sharp changes in $\mathbf{N_e}$, and the signal should increasingly overwhelm regularization. This automatic hyperparameter selection scheme based on an intuitive heuristic is not foolproof. We always recommend examining how the estimated $\mathbf{N_e}$ fits the observed IBD segment distribution to explore whether over-regularization occurs (see Fig.S2 for one example).

### Supp. Note S2 Estimating IBD calling Error Parameters for Empirical aDNA Data

As described in the main text, we consider three types of IBD detection errors: false positives, imperfect power, and length bias. The following subsections describe how we estimated these three error types from simulations.

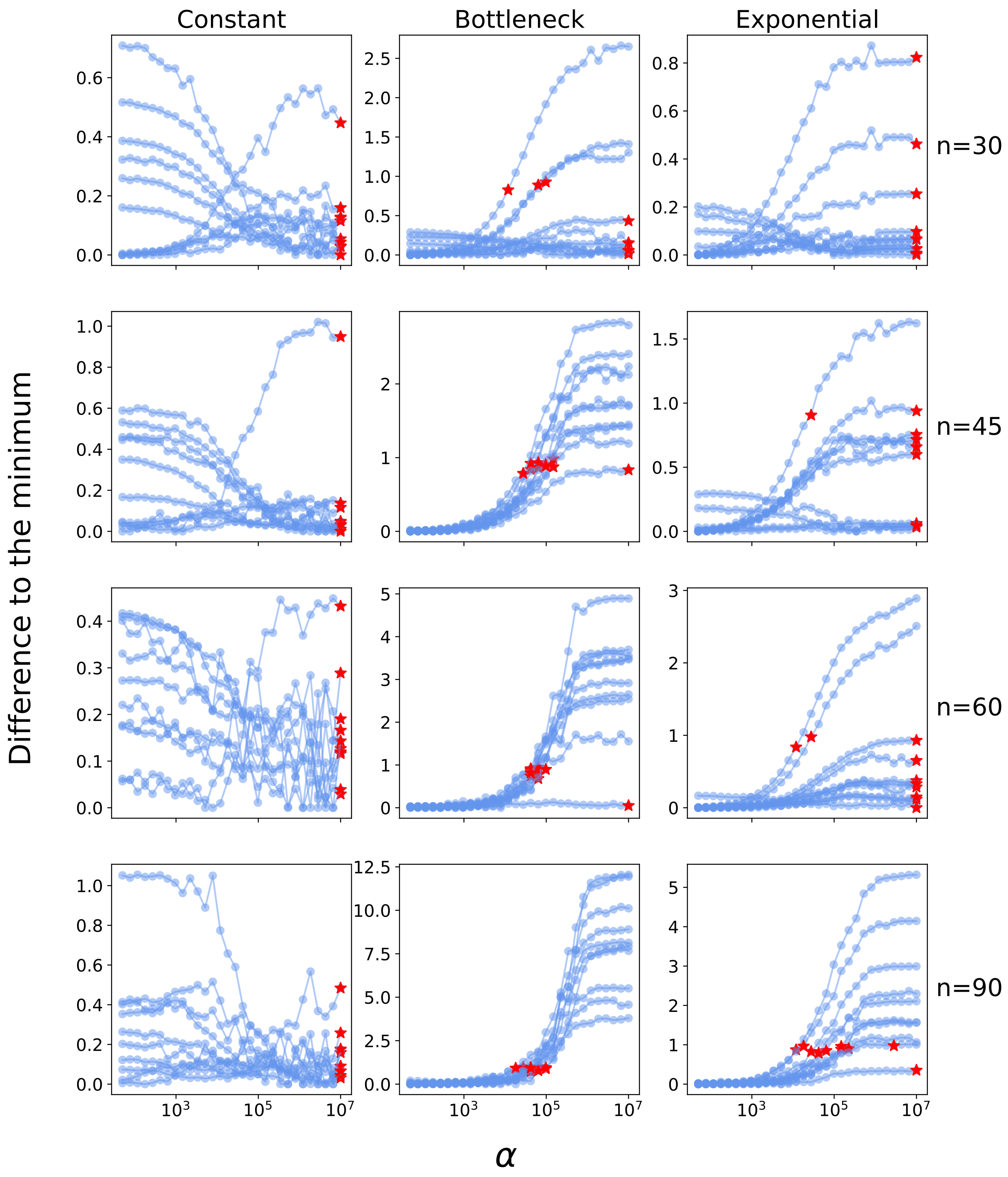


**Figure S1: Effects of $\alpha$ on deviance statistics.** We visualize deviance statistics $D(\alpha)$ for varying $\alpha$ values (i.e. strength of regularization). The y-axis is the difference to the minimum deviance statistics (over the search space of $\alpha$). We show the results using a single sampling point at $t=0$, for various sample sizes (rows). Ten independent simulated replicates are depicted for each scenario (blue curves). We indicate $\alpha^*$, the "optimal" $\alpha$ we chose for each replicate (red star). We observed that models with multiple sampling time points have qualitatively similar behaviors (results not shown).

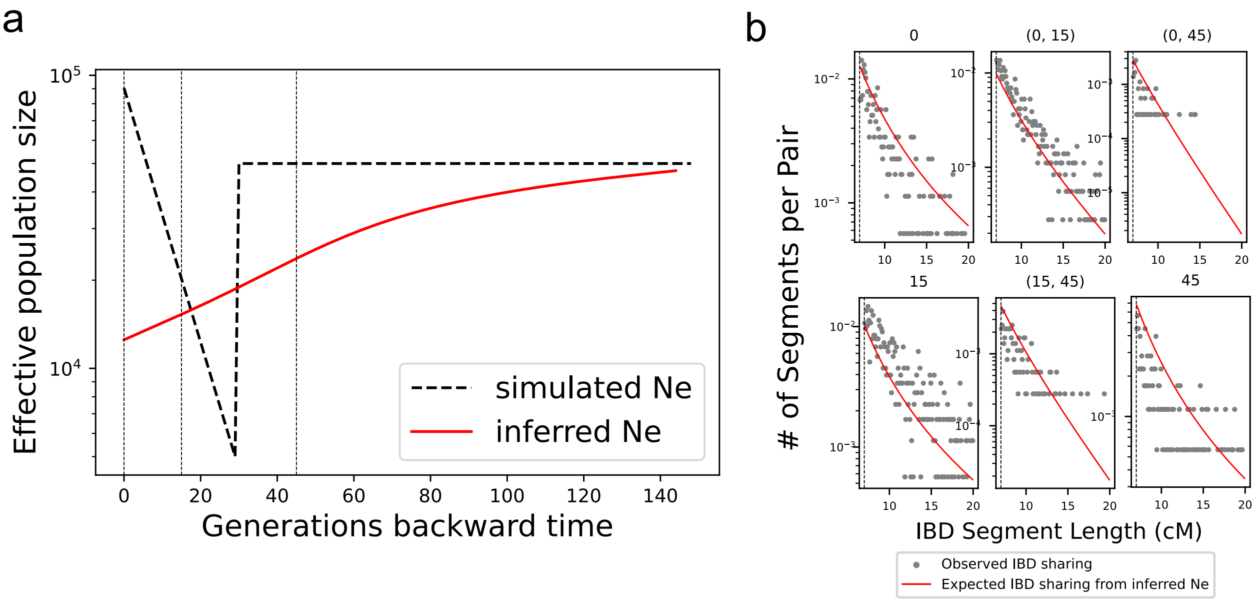


**Figure S2: An example of over-regularization.** We depict an example of over-regularized estimates of the bottleneck demography (when manually fixing $\alpha$=10,000,000). **a** Inferred $\mathbf{N_e}$ (red) vs. simulated $\mathbf{N_e}$ (dashed black line).The bottleneck occurs at $t$=30, and the three sampling times are $t$=0,15,45 (vertical dashed black line). **b** Comparison of empirical IBD to that from model fit. The sub-figure title indicates the sampling time (in generations). For example, 0 indicates that this subplot shows the empirical (gray dots) and predicted (red line) IBD within the first sample set sampled at $t$=0. (0,15) indicates that this subplot depicts the empirical and predicted IBD between the first and second sample set, sampled at $t$=0,$t$=15, respectively. We note that the IBD sharing expected from the estimated $\mathbf{N_e}$ does not fit well with the empirical IBD within the $t$=15 samples and across the $t$=0,$t$=15 and $t$=15,$t$=45 pairs. Because the model is over-regularized, the estimated $N_e$ around the bottleneck period gets over-smoothed, and the model cannot recapitulate the large population size before the bottleneck. The former leads to underestimated IBD sharing between the $t$=0 and $t$=15 sample clusters, and the latter leads to overestimated IBD sharing between $t$=15 and $t$=45 sample clusters.

#### Supp. Note S2.1 Estimating False Positive Rates

We empirically estimated the false positive rate for the average coverage of samples. The estimation procedure is the same as in Supplementary Note 7 in Ringbauer et al. [2023]. Briefly, 11 samples (Supplementary Table 1) with no IBD were selected and downsampled to the desired coverage (in this case, the average coverage of samples used in our empirical data analysis). Then the false positive rates at various IBD lengths were estimated from downsampled data, using 100 independently downsampled replicates.

#### Supp. Note S2.2 Estimating IBD Length Bias

We define IBD length bias as the difference between the inferred and actual IBD length. To correctly estimate this bias, we need simulations where the precise boundary of ground truth IBD is known, and the data should be qualitatively similar to empirical aDNA data. Toward this end, we started with empirical aDNA BAM files on chromosome 3 and copied in IBD segments at known locations. We first subset reads by genomic regions they aligned to and then merge reads from different genomic regions and individuals to form synthetic pairs of individuals with IBD shared at defined genomic positions. In particular, we utilize the fact that parent-offspring pairs naturally share IBD along their whole genome. Therefore, to simulate a pair of individuals with one IBD of length $l$cM, we first randomly select a $l$cM along the chromosome. The first simulated individual takes reads aligned to the selected region from the BAM file of the parent, and the second individual from the BAM file of the child. For the rest of the genomic regions, the two simulated individuals take reads from the BAM files of two genomes without IBD. We illustrate this approach graphically in Fig.S3a.

To simulate IBD sharing in WGS-like data, we used the WGS BAM files of a high coverage father-son pair (I3950, I3949) published in Wohns et al. [2022] for the IBD region. This father-son pair is associated with Afanasievo culture and was dated to 2879-2632 calBCE and 2844-2496 calBCE, respectively. To simulate IBD sharing in 1240k-like data, we used the 1240k BAM files of a mother-son pair (GRG080, GRG041) published in Rivollat et al. [2023]. The mother-son pair originates from a Neolithic burial site (Gurgy 'les Noisats') in present-day France from 4850–4500 BC. For the non-IBD region, we used I3255 (2139-1947calBCE, England_Bellbeaker, [Olalde et al., 2018]) and I2105 (3300-2800BCE, Ukraine_EBA_Yamnaya, [Mathieson et al., 2018]) for each of the two individuals in a simulated pair. No IBD sharing $>$ 5cM is detectable in this pair - as the two individuals are separated by ca. 1000 years. The I3255 and I2105 were initially published with 1240k-capture adNA data and were later also WGS-sequenced to high-coverage (data publicly available at `https://reich.hms.harvard.edu/ancient-genome-diversity-project`). We used the respective data types for our WGS and 1240k simulations.

For the Corded ware samples used in our empirical analysis (we performed analogous simulations for the UK data), the average coverage for 1240k data is 1.65x (calculated on the 1240k SNP set), and that for WGS data is 1.1x. Therefore, we simulated IBD of 8cM, 12cM, 16cM, and 20cM with coverage and data type corresponding to the empirical data (Fig.S3b). To calculate length bias, for each simulated groundtruth IBD, we recorded inferred IBD segments that cover at least half of the groundtruth segment (therefore, there can only be one unique inferred segment satisfying this condition, if it exists at all). The length bias is the length difference between the inferred and the groundtruth segments. If no such inferred segment exists, then this ground truth segment is considered not inferred, and no length bias is recorded.

We found that the length bias is asymmetrical (Fig.S3b,c). Most segments are inferred to be within 2cM longer than their groundtruth lengths. However, a much smaller albeit non-negligible proportion of segments are broken apart due to errors in the imputed genotypes. In addition, we found that length bias is similar across different IBD lengths (Fig.S3b). Therefore, we modeled length bias across different groundtruth IBD lengths with the same model for simplicity. We also found that length bias is overall smaller for 1.1x WGS data compared to 1.6x 1240k data, consistent with the results of Ringbauer et al. [2023], which finds that in terms of IBD calling performance WGS data is similar to 1240k data with approximately 3x more coverage. The length bias distribution for 1.1x WGS appears similar to that of 1.65x 1240k data (Fig.S3b,c); therefore, we estimated a single-length bias distribution using the combined simulated WGS and 1240k data. We simulated 12cM IBD with 500 independent replicates for each 1240k and WGS data. Then, we fitted a single density estimation (KDE) with Gaussian kernels to estimate the length bias distribution (shown by red line in Fig.S3c).

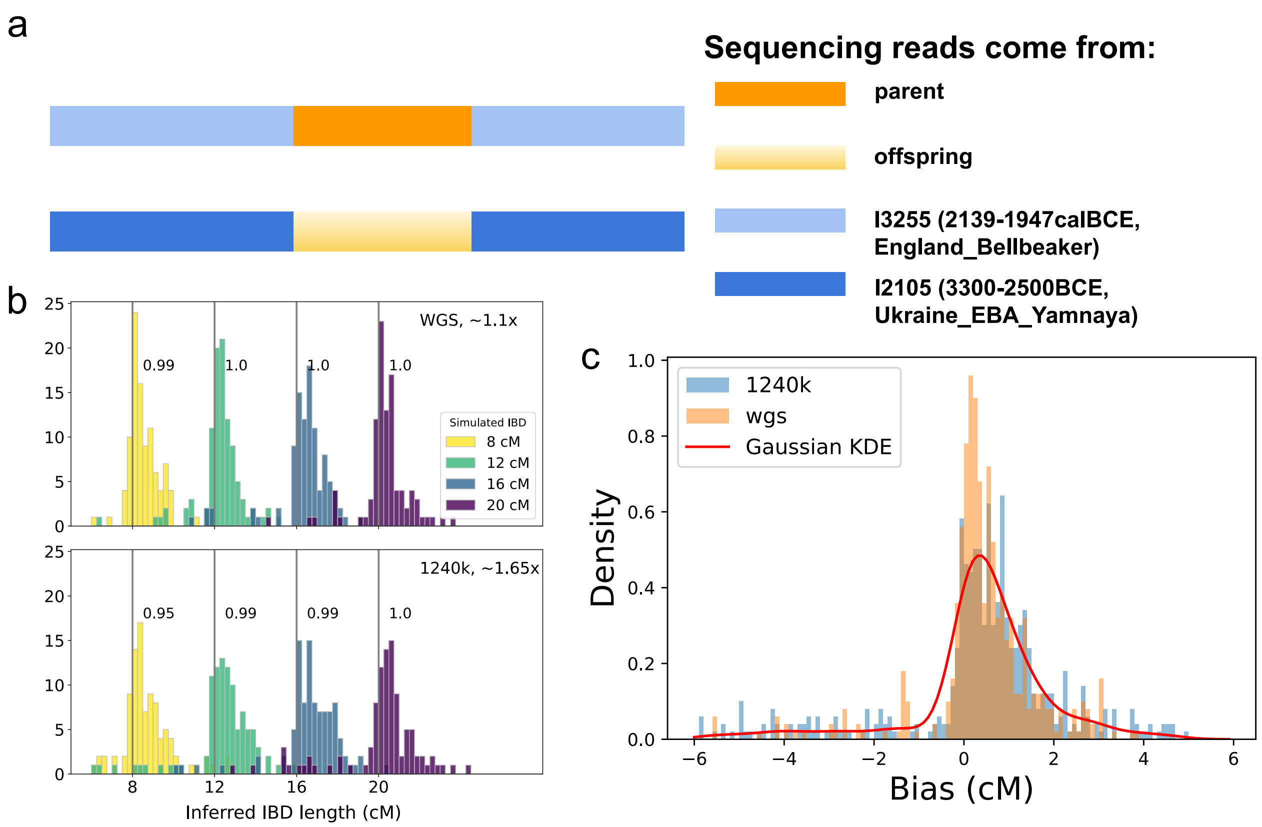


**Figure S3: Estimating IBD length bias from simulated data.** **a** A schematic showing our procedure of mixing sequencing reads aligned to different genomic regions to simulate IBD. **b** Histogram of inferred segment lengths on simulated IBD with 1.1x WGS data and 1.65x 1240k data. For each data type and IBD length, we simulated 100 independent replicates and then plotted the histogram of inferred IBD length. The number next to the vertical line indicates the recall for the corresponding segment length class. **c** Distribution of length bias of simulated 12cM groundtruth IBD for 1240k (shown in blue) and WGS data (shown in orange). We simulated 400 independent replicates in addition to that shown in **b** to fit the KDE. A red line shows the fitted Gaussian KDE.

#### Supp. Note S2.3 Estimating Recall

We use the same simulation scheme explained in the previous section (Supp. Note Supp. Note S2.2) to estimate recall. We simulated groundtruth IBD of 2cM,3cM,4cM,5cM,6cM,

7cM,8cM,9cM,10cM,11cM,12cM,16cM,20cM. Similar to Supp. Note S2.2, a groundtruth IBD segment is considered to be inferred if an inferred segment covers at least half of the true IBD segment. Because in calling IBD in empirical aDNA data we used length threshold of 6cM, here we apply the same length threshold of 6cM, regardless of the groundtruth segment length. Consequently, the recall for true segments less than 6cM is very low. We found that the recall for 1.65x 1240k data is similar to that of 1.1x WGS data (Fig.S4). In the error model for our empirical data analysis, we use the average of the two. We linearly interpolated recall for segment lengths between the simulated IBD lengths.

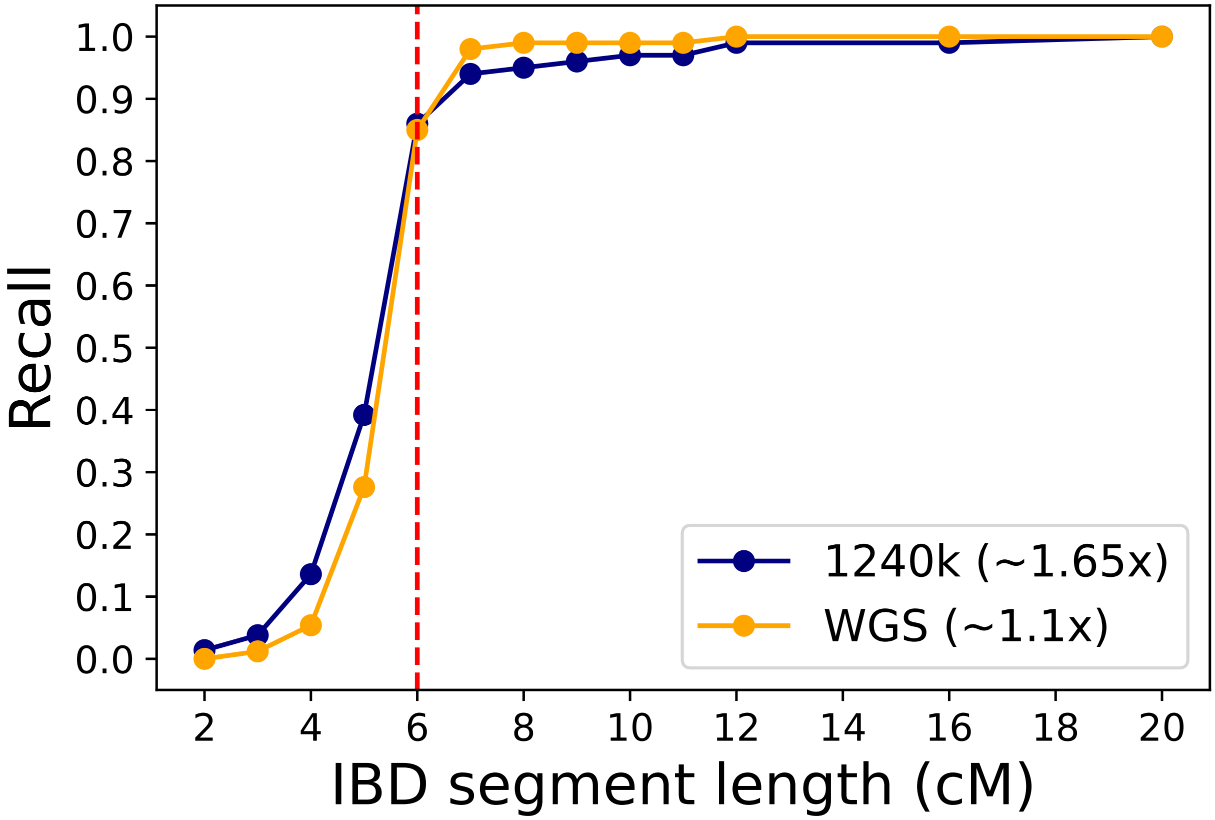


**Figure S4: Recall as a function of segment length.** We simulated groundtruth IBD of various lengths (see Fig.S3) and recorded ANCIBD's recall for 1240k (in navy) and WGS (in orange) data. We simulated 500 independent replicates for groundtruth segment less than 6cM and 100 replicates for the longer length classes.

#### Supp. Note S3 Sampling Time Uncertainty

Unlike contemporary DNA samples, aDNA samples typically do not have an exact date. Instead, they are dated either by radiocarbon dating or archaeological context, giving a plausible sample time range. This time interval can be broad, such as during the Hallstatt plateau that spans 400 years [Stuiver and Pearson, 1986]. Therefore, we describe how we can incorporate such sampling time uncertainty into our model and investigate how time uncertainty biases our estimates.

We assume all samples within a sample set $S_i$ have the same time range. This may not be entirely accurate in practice; however, we consider this a reasonable modeling assumption to keep the model tractable while accommodating some of the complexities of empirical data. Each sampling time point $t_i$ is associated with a radius $r_i$ such that all samples in $S_i$ are assumed to be uniformly sampled from the time range $[t_i - r_i, t_i + r_i]$. Our model and implementation do not require the interval to be centered around $t_i$, but we assume the time interval is symmetrical around $t_i$ for notational simplicity. In practice, the radius $r_i$ could be determined from 95.4% CI of calibrated radiocarbon age. Consider two sampling clusters $S_1, S_2$, each with time range $[t_1 - r_1, t_1 + r_1]$ and $[t_2 - r_2, t_2 + r_2]$. Denote the likelihood of observed IBD segments between $S_1, S_2$ with fixed sampling time $t_1^{'}, t_2^{'}$ as $\mathcal{L}(S_1, S_2 | \mathcal{N}, t_1^{'}, t_2^{'})$, which can be calculated as described in Methods, we can then integrate the likelihood over the time range as follows,

$$\begin{aligned}\mathcal{L}(S_1,S_2|\mathcal{N}) &= \int_{t_1-r_1}^{t_1+r_1}\int_{t_2-r_2}^{t_2+r_2}\mathcal{L}(S_1,S_2|\mathcal{N},t_1^{'},t_2^{'})P(t_1^{'},t_2^{'})dt_1^{'}dt_2^{'} \\ &= \frac{1}{4r_1r_2}\sum_{t_1^{'}=t_1-r_1}^{t_1+r_1}\sum_{t_2^{'}=t_2-r_2}^{t_2+r_2}\mathcal{L}(S_1,S_2|\mathcal{N},t_1^{'},t_2^{'})\end{aligned} \quad (1)$$

To explore the effect of having samples originating from different time points but modeled as contemporaneous, we simulated the constant and bottleneck model with samples taken uniformly from a time interval (specified by radius $r$) centered at specified time points. For example, for radius $r$ and time point $t$, samples are uniformly taken from the time interval from $t - r$ to $t + r$. We found that when $r$ is large (e.g., $r = 10$), not accounting for time heterogeneity leads to overestimates of $N_e$. Although the effect is moderate, this bias aligns with the findings from Fournier et al. [2023]. The correction procedure described in Eq.1 alleviates this upward bias (Fig.S5). However, as expected, this correction procedure runs $r^2$ times slower because of the double summation in Eq.1.

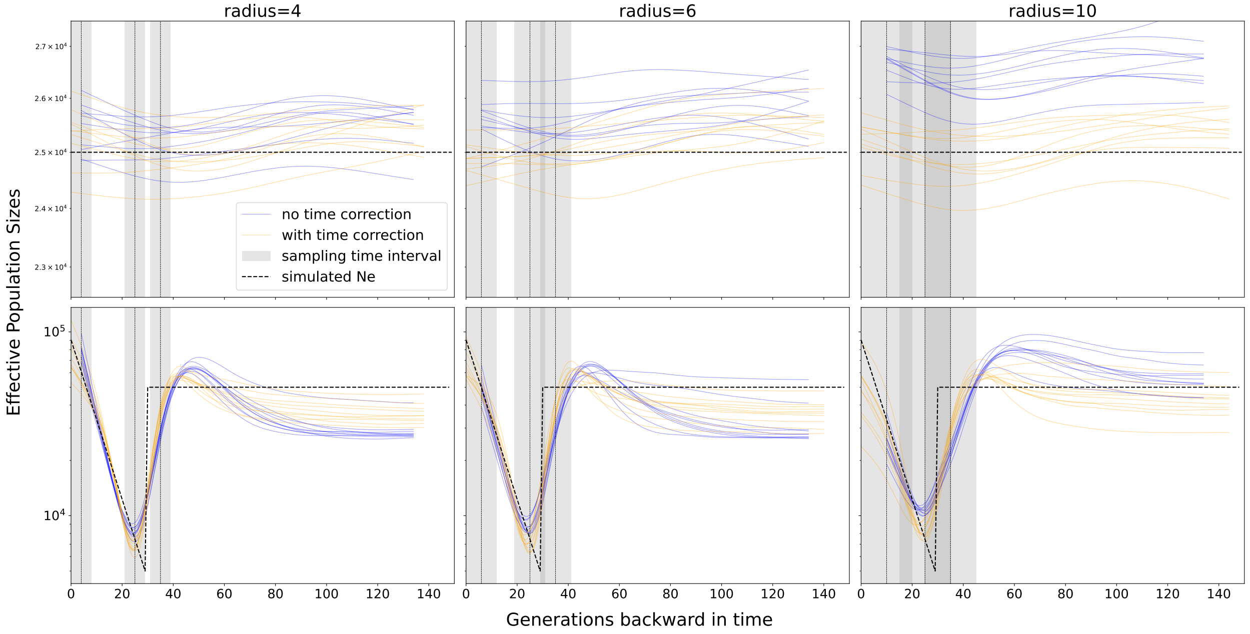


**Figure S5: Simulated constant and bottleneck demography with samples drawn from intervals of various widths** This plot shows the inferred $\mathbf{N_e}$ of simulated constant/bottleneck demography with samples uniformly drawn from intervals of width 8,12,20 generations (with and without the time heterogeneity correction in Eq.1). The vertical dashed lines indicate the mean sample ages for each of the three sampling points, and the grey-shaded area represents the time interval over which samples are drawn. The inferred $\mathbf{N_e}$ without Eq.1 correction does not start at $t = 0$ because the model assumes that the first sampling point is at the mean age of the most recent set of samples.

### Supp. Note S4 Additional Supplementary Figures

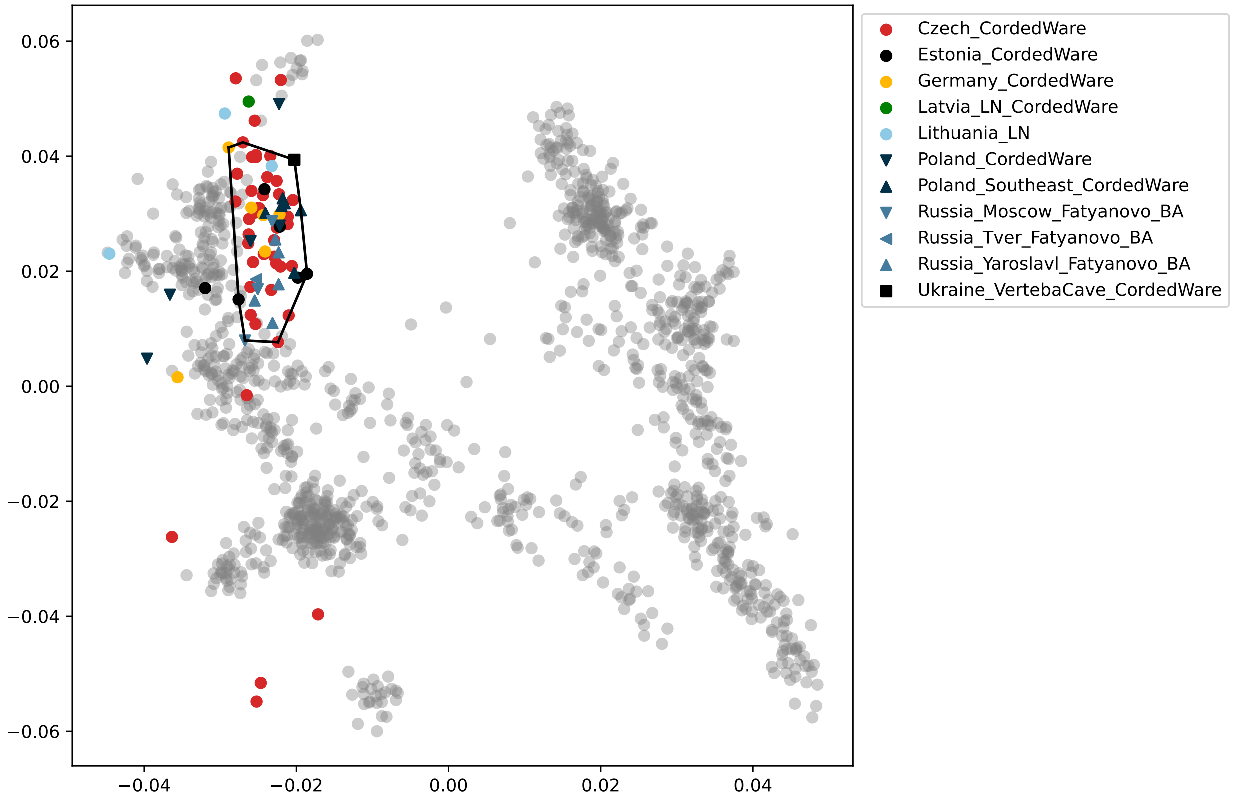


**Figure S6: PCA of published Corded Ware samples.** Principle Component Analysis (PCA) of published Corded Ware samples passing coverage requirement for HapNe-LD (having at least 300k 1240k SNPs covered, as recommended in Fournier et al. [2023]). The gray dots are the typical West-Eurasian Human Origins samples widely used as a PCA reference dataset for aDNA studies [Mallick et al., 2024]. The black boundary indicates the CW genomes we used to infer population size trajectory. We exclude individuals falling outside the black boundary as ancestry outliers. For HapNe-LD analysis, we used all samples with at least 300k 1240k SNPs covered, as recommended Fournier et al. [2023]. For ANCIBD analysis, we included all samples with at least 600k 1240k SNPs covered, a subset of samples shown in this PCA plot.

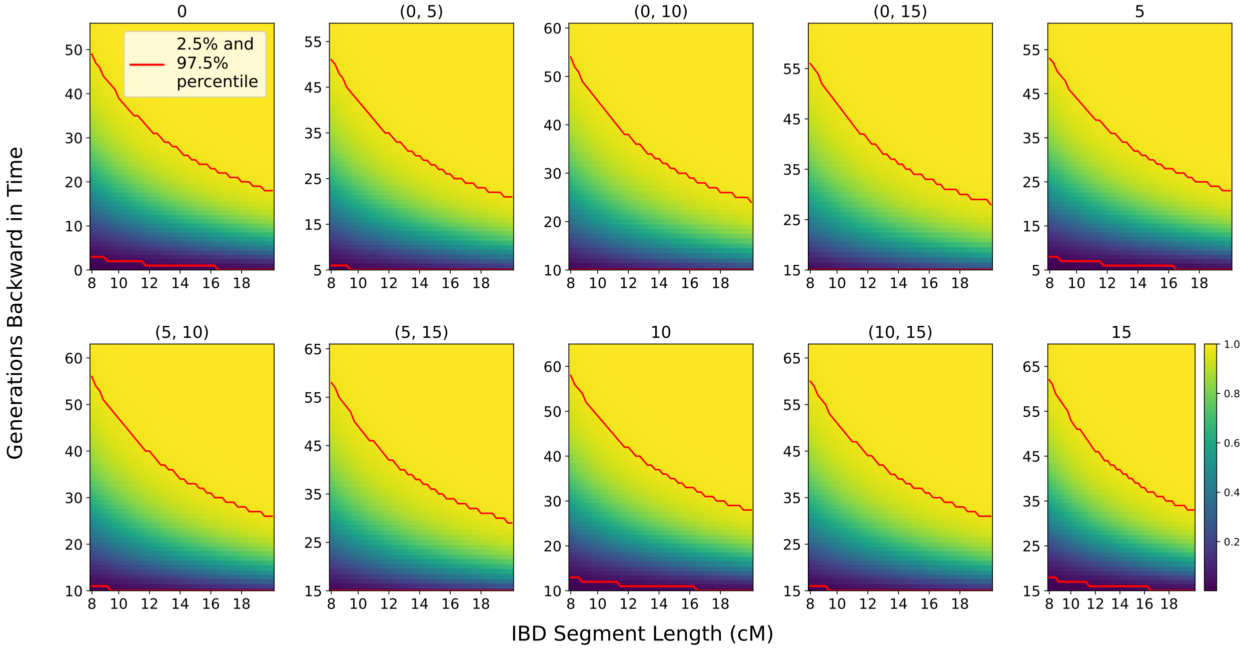


**Figure S7: Cumulative Density of TMRCA for IBD segments of various lengths.** Cumulative density plot of TMRCA of IBD segments of various lengths (x-Axis) calculated when using the inferred CW demography from IBD-sharing in CW individuals (see Fig. 5 in the main text). The two red lines depict the 2.5% and 97.5% percentile.

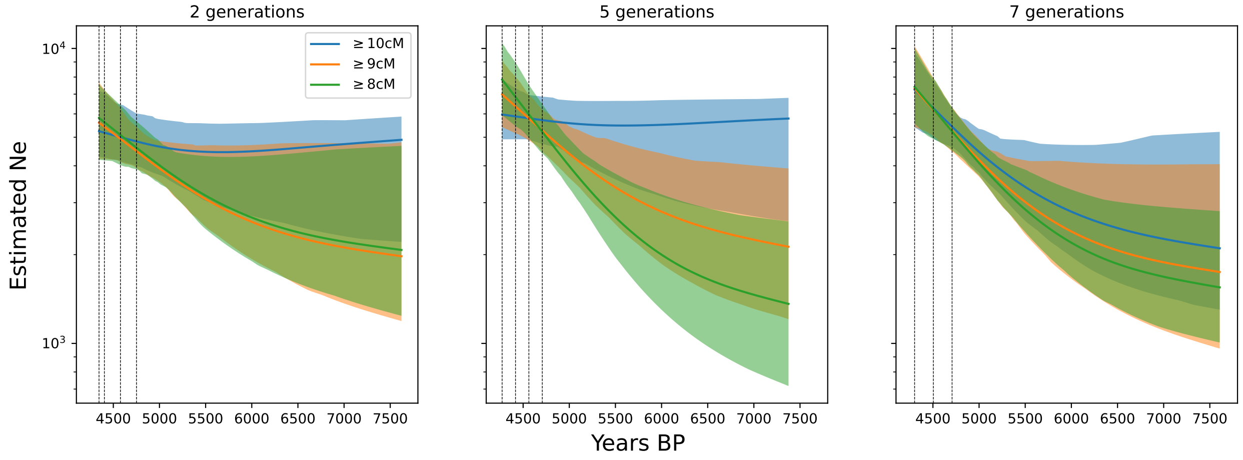


**Figure S8: Inferred CW $N_e$ trajectories across different levels of sample binning.** We tested different levels of sample binning (e.g., grouping samples whose median radiocarbon date or archaeological context date are within 2,5,7 generations of one another) and inferred Ne trajectories. As a convention, we took 29 years as the generation time.

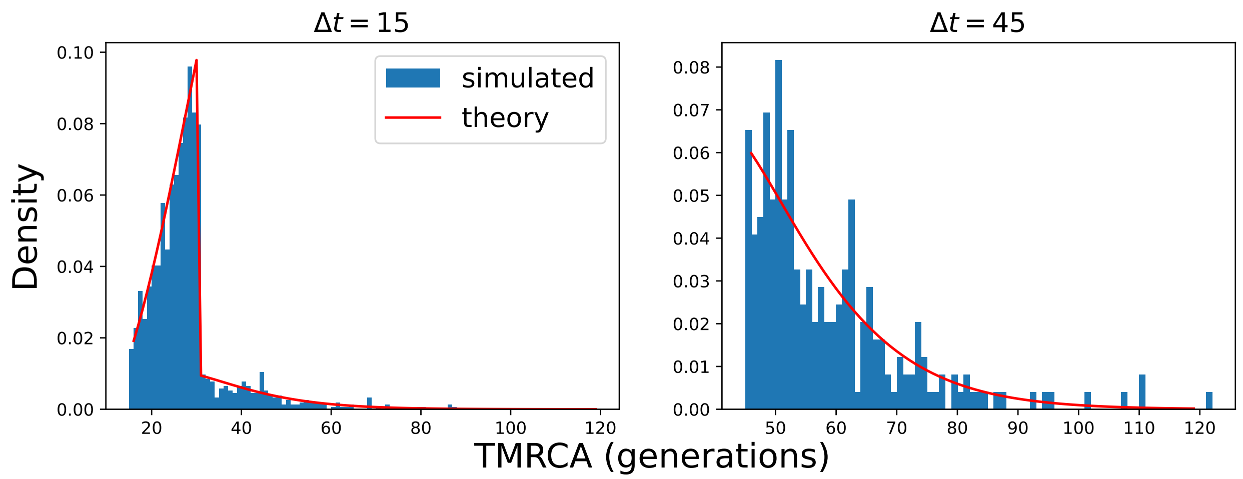


**Figure S9: Comparing TMRCA of simulated IBD segments with analytical predictions.** We simulated IBD segments under the bottleneck demography as in Fig.2b, except that we scaled the effective population size by a factor of 0.1 to observe a sufficient number of IBD segments from 10,000 replicates to study the statistical properties of their TMRCA. We recorded the TMRCA of IBD segments shared between two haplotypes, one sampled at $t=0$, and the other at $t=15$ (left) or $t=45$ (right). We plotted the TMRCA distribution of simulated IBD segments with lengths between 5.75cM and 6.25cM as a histogram (in blue) and superimposed posterior TMRCA distribution theoretically calculated as described in Methods (in red), showing that the theoretical calculation matches the empirical simulations.

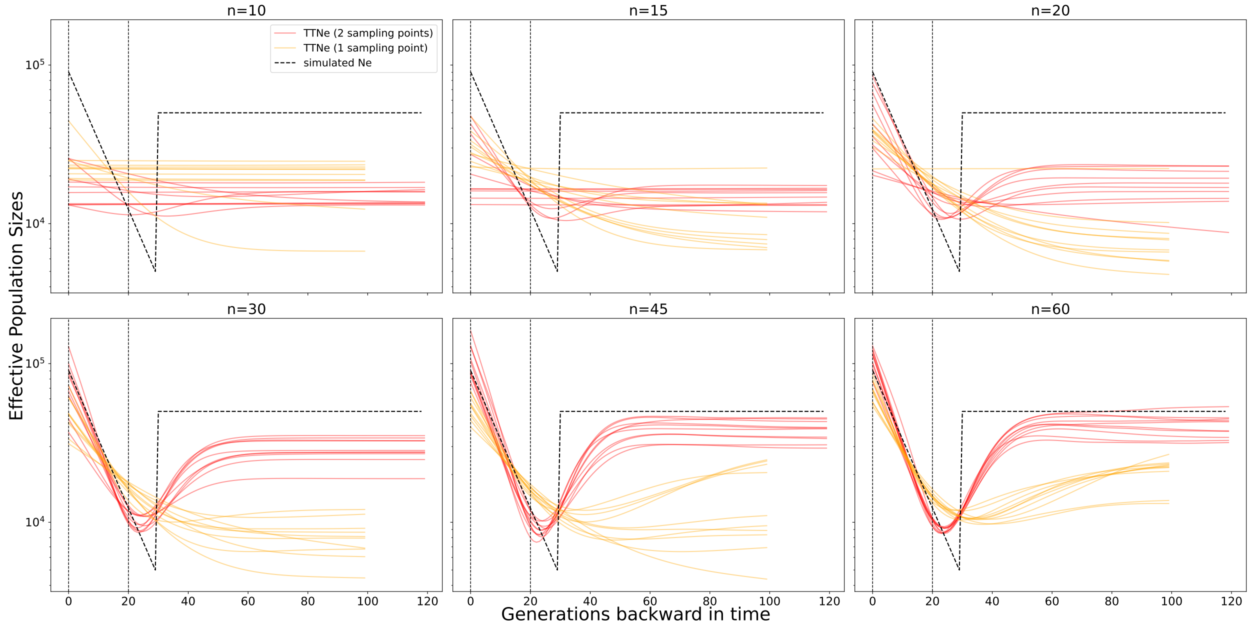


**Figure S10: Inferring $N_e$ in the bottleneck demography with two sampling points after a bottleneck event.** Results of inferred Ne with two sampling points in the bottleneck demography (in red). The vertical dashed lines indicate the two sampling points. The subtitle indicates the sample size taken at each time point. As a baseline comparison, we also plot the results of inferred Ne using only contemporaneous samples (visualized in orange). The baseline has $3n$ samples at $t=0$. Although a comparison with $2n$ samples at $t=0$ is fairer, we did not conduct additional simulation with $2n$ samples as this does not seem necessary.

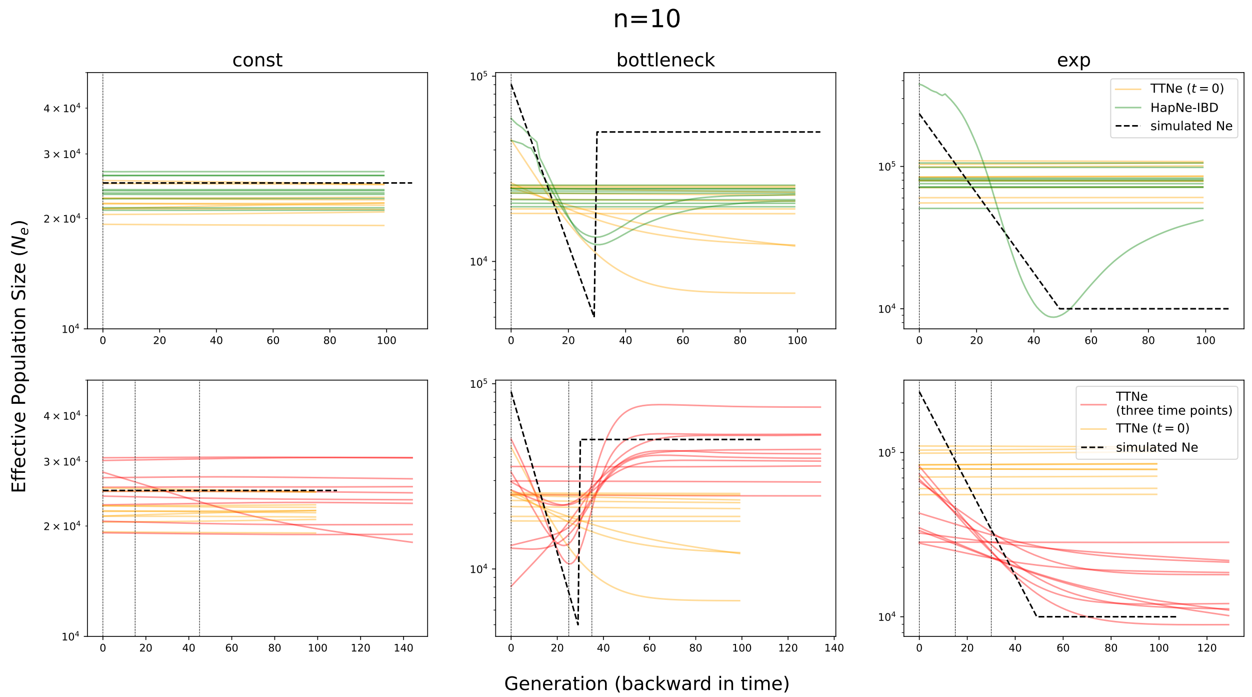


**Figure S11: Performance of TTNE in various simulated demographic scenarios with** $n = 10$**.** Same as Fig. 3 in the main article but with $n = 10$ at each sampled time point for models with multiple sampling points or $n = 30$ for models with contemporaneous samples only.

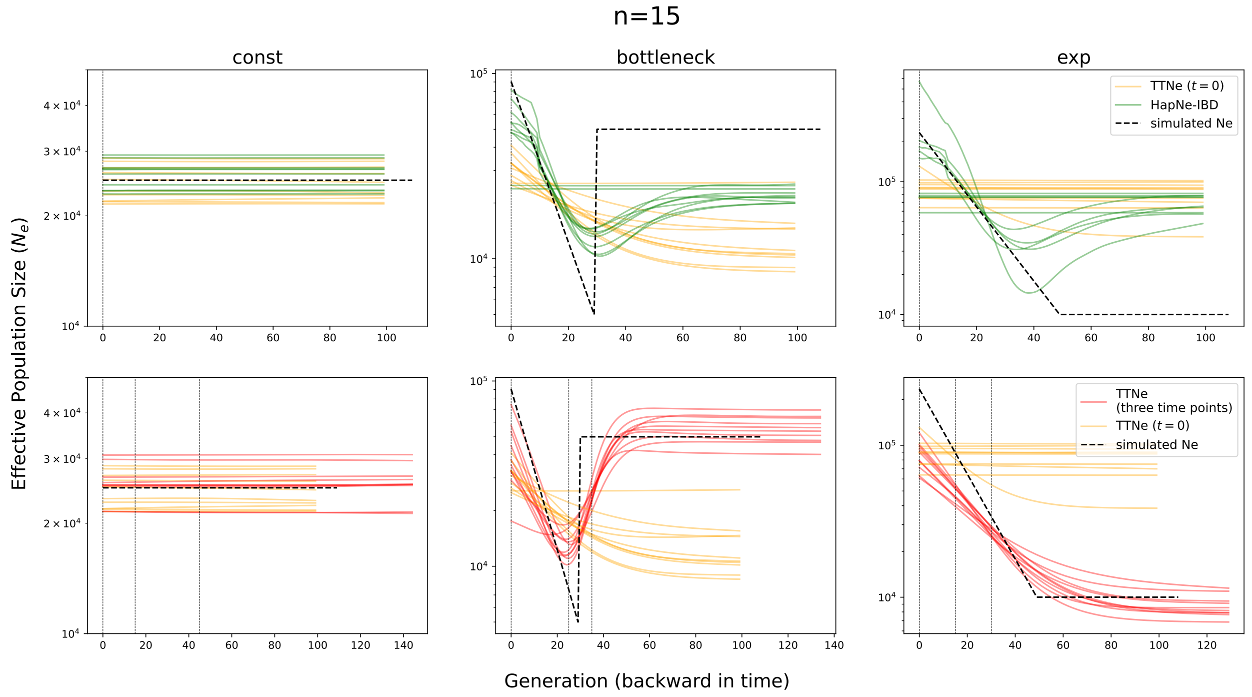


**Figure S12: Performance of TTNE in various simulated demographic scenarios with** $n = 15$**.** Same as Fig. 3 in the main article but with $n = 15$ at each sampling time point for models with multiple sampling points or $n = 45$ for models with contemporaneous samples only.

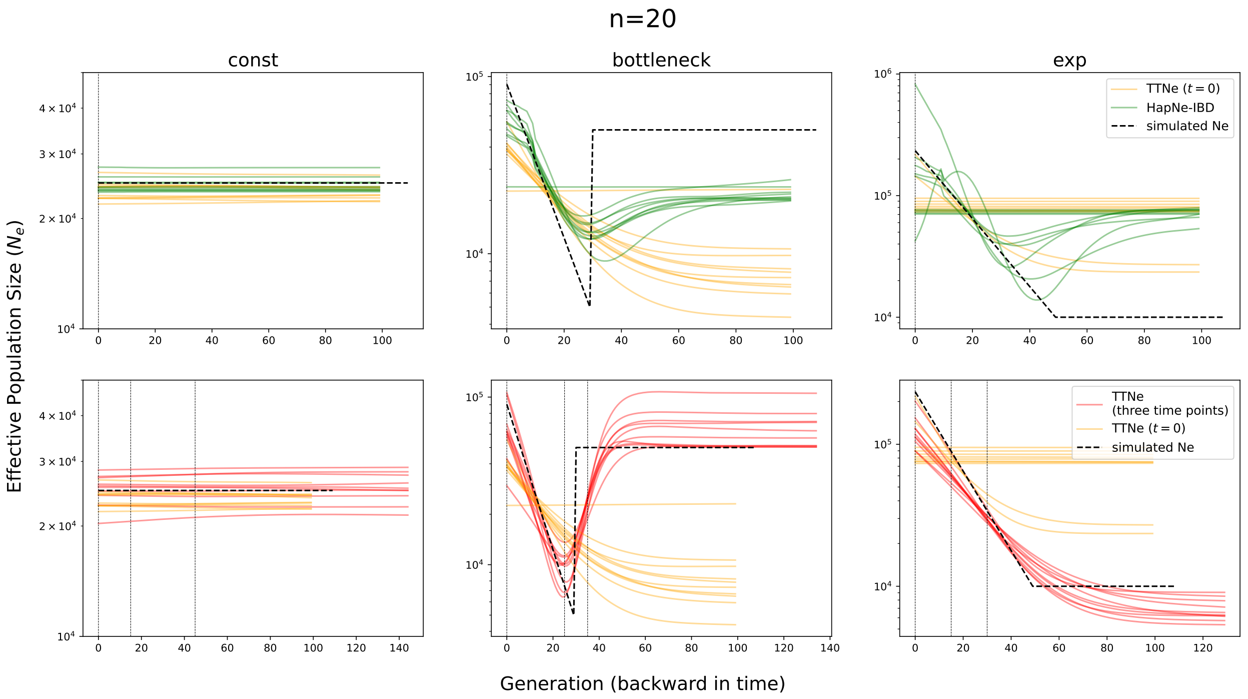


**Figure S13: Performance of TTNE in various simulated demographic scenarios with** $n\!=\!20$**.** Same as Fig.3 in the main article but with $n\!=\!20$ at each sampled time point for models with multiple sampling points or $n\!=\!60$ for models with contemporaneous samples only.

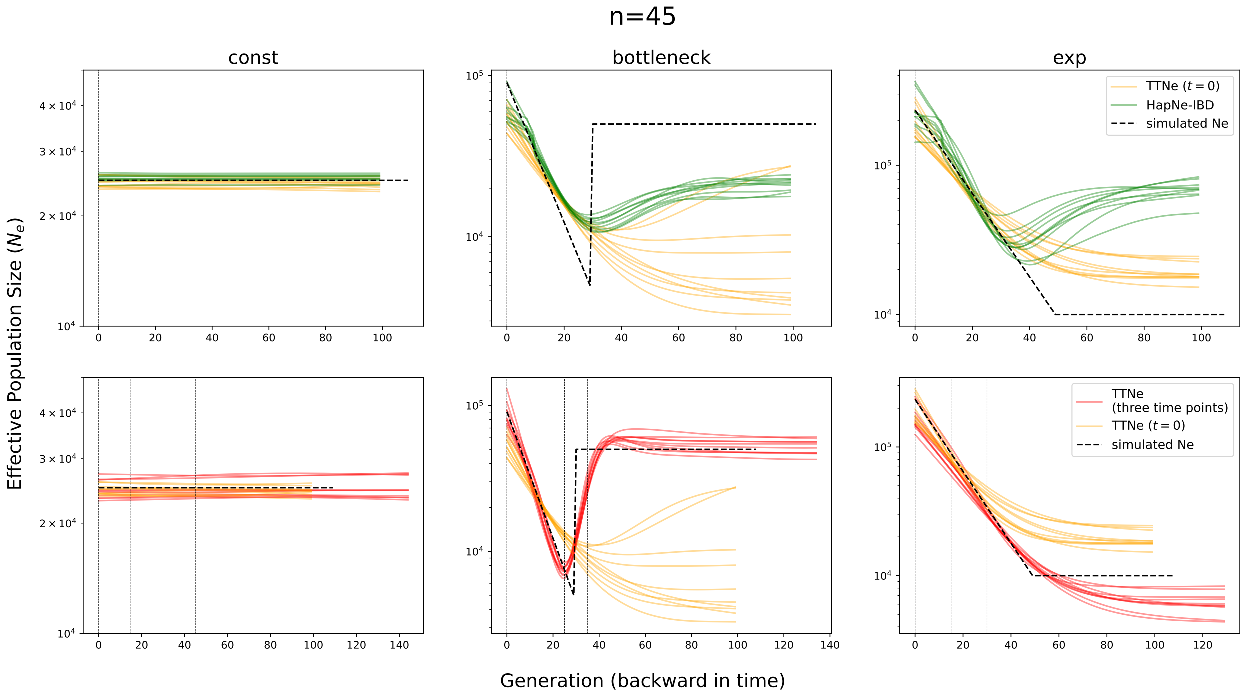


**Figure S14: Performance of TTNE in various simulated demographic scenarios with** $n\!=\!45$**.** Same as Fig.3 in the main article but with $n\!=\!40$ at each sampled time point for models with multiple sampling points or $n\!=\!135$ for models with contemporaneous samples only.

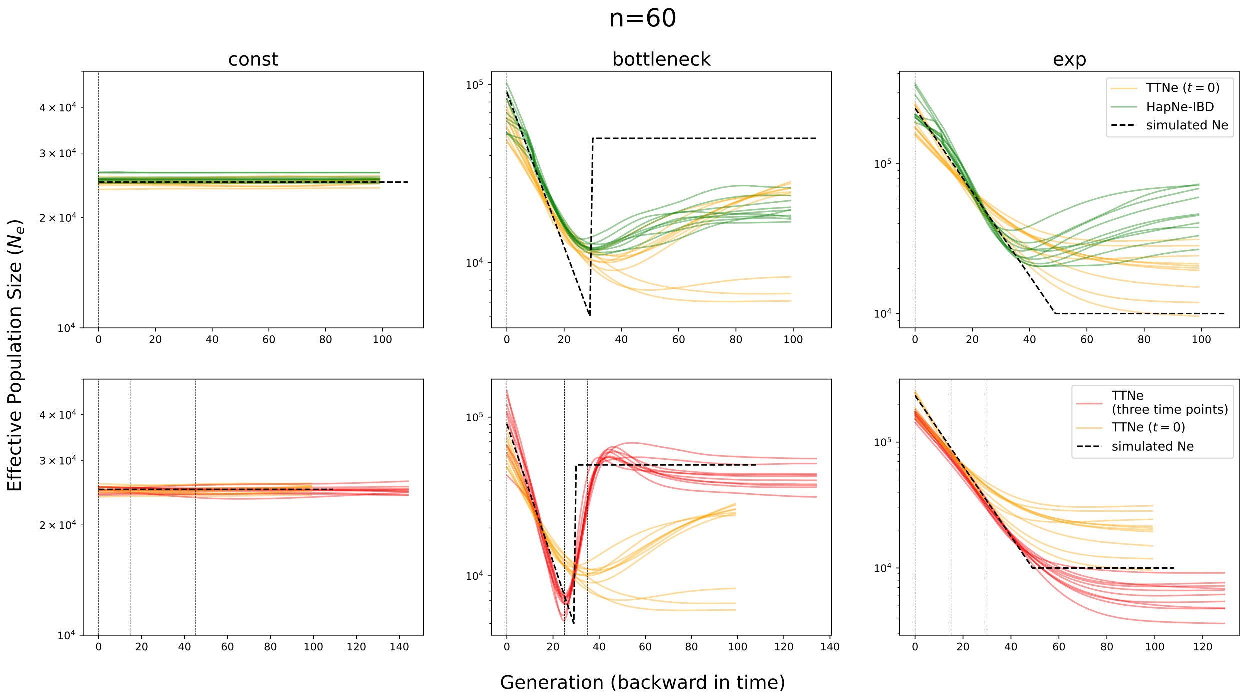


**Figure S15: Performance of TTNE in various simulated demographic scenarios with $n=60$.** Same as Fig.3 in the main article but with $n=60$ at each sampled time point for models with multiple sampling points or $n=180$ for models with contemporaneous samples only.

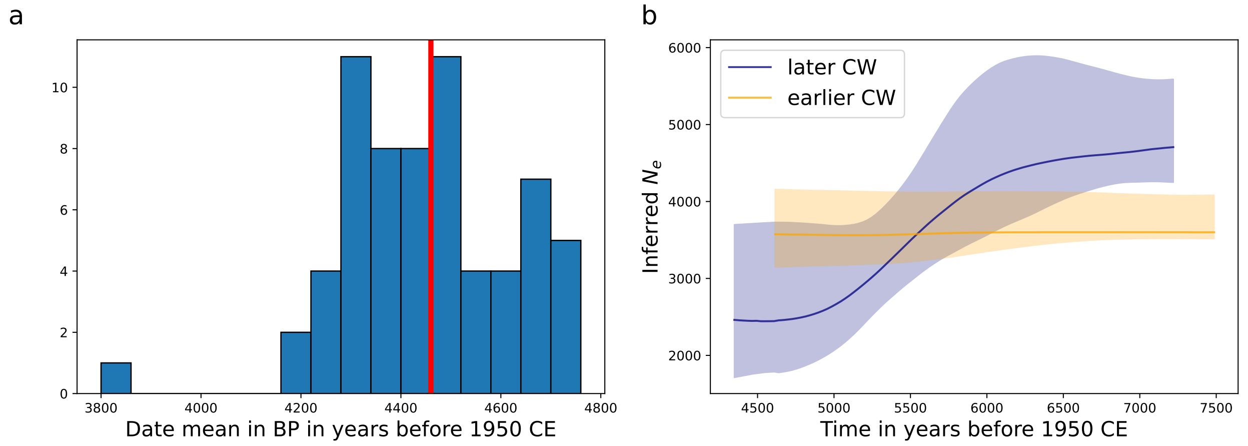


**Figure S16: Inferring Corded Ware Ne using hapNe-LD a** We plotted the histogram of the mean date (in BP in years before 1950CE) of individuals falling inside the main Corded Ware cluster (as shown in Fig.S6). We divided them into two groups (indicated by the thick red vertical bar), the earlier CW cluster consists of individuals dated before 4460BP, and the later CW cluster consists of individuals dated after 4460BP (but excluding VERT113B, an individual dated to 3823BP), and applied hapNe-LD separately to each of the two groups. **b** Inferred $N_e$ of the earlier (in orange) and later CW (in navy) using hapNe-LD. For both groups, hapNe-LD warned about cross-chromosome LD, indicating that recent admixture may substantially bias the results.

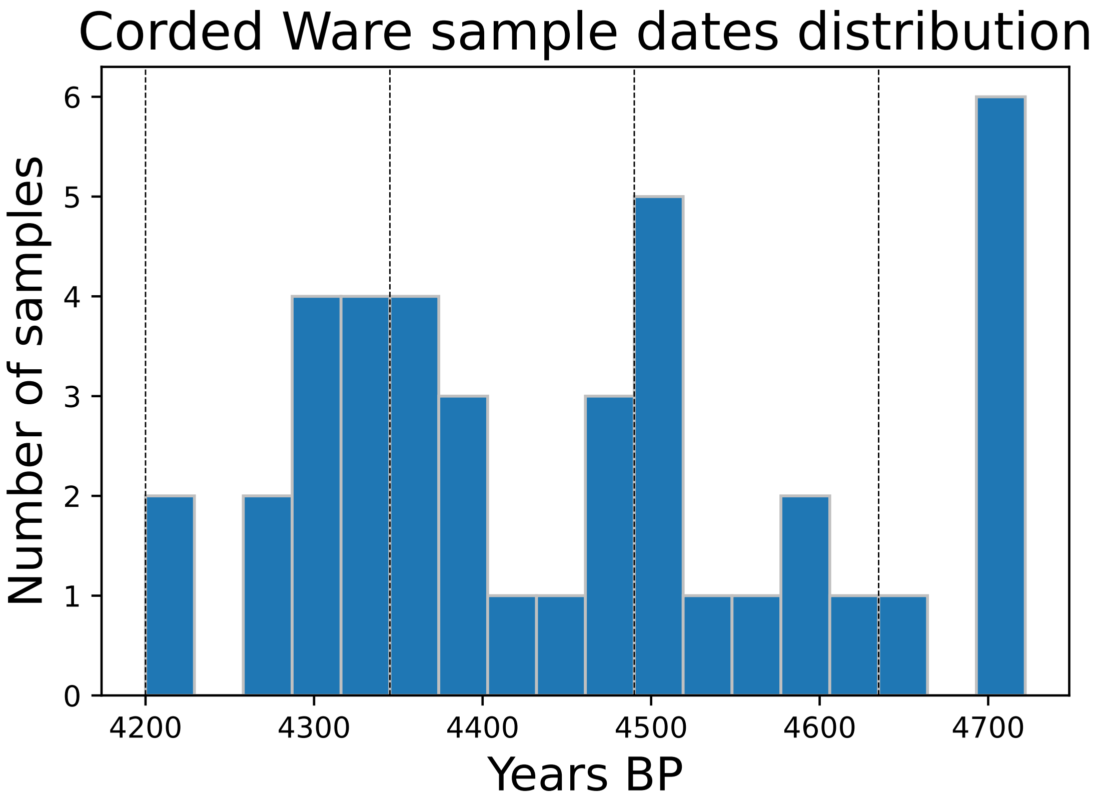


**Figure S17: Date distribution of CW samples used for IBD analysis.** Date distribution of CW samples used for IBD analysis and the grouping of temporally close samples. The dashed black vertical lines indicate the date delimiter of each temporal group. Each group spans 145 years (or five generations, assuming a generation time of 29 years).

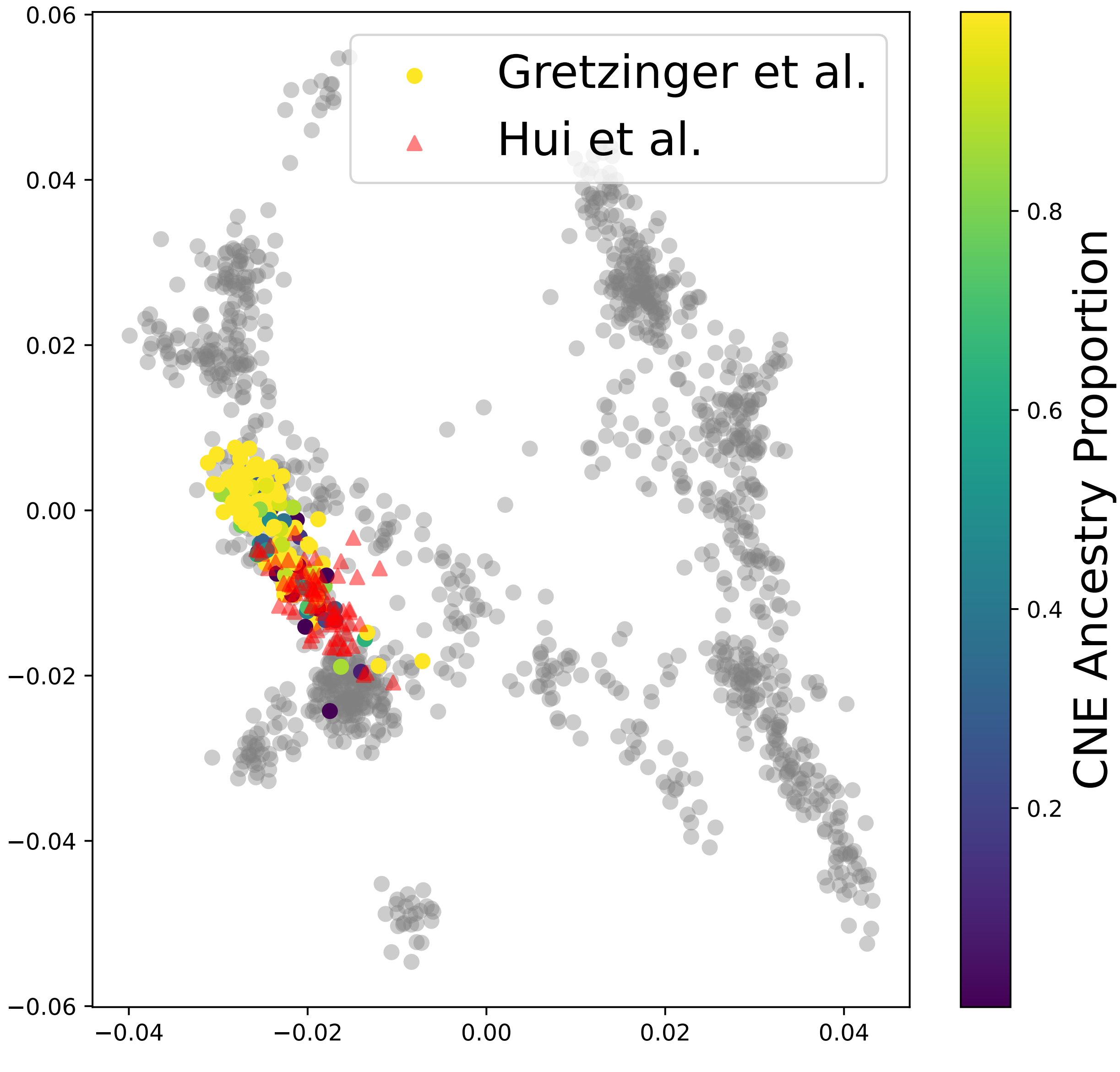


**Figure S18: PCA of samples from the British Isle.** PCA of samples from Gretzinger et al. [2022] and Hui et al. [2024]. For the samples from Gretzinger et al. [2022], we color-coded each individual according to its CNE (continental northern European) ancestry proportion. The CNE ancestry proportion was reported in Supplementary Table S3.7 in Gretzinger et al. [2022] by performing supervised admixture with source populations CNE and WBI (Western British and Irish).
